# Rapid, scalable, combinatorial genome engineering by Marker-less Enrichment and Recombination of Genetically Engineered loci (MERGE)

**DOI:** 10.1101/2022.06.17.496490

**Authors:** Mudabir Abdullah, Brittany M. Greco, Jon M. Laurent, Michelle Vandeloo, Edward M. Marcotte, Aashiq H. Kachroo

## Abstract

Large-scale genome engineering in yeast is feasible primarily due to prodigious homology-directed DNA repair (HDR), a plethora of genetic tools, and simple conversion between haploid and diploid forms. However, a major challenge to rationally building multi-gene processes in yeast arises due to the combinatorics of combining all of the individual edits into the same strain. Here, we present an approach for scalable, precise, multi-site genome editing that combines all edits into a single strain without the need for selection markers by using CRISPR-Cas9 and gene drives. First, we show that engineered loci become resistant to the corresponding CRISPR reagent, allowing the enrichment of distinct genotypes. Next, we demonstrate a highly efficient gene drive that selectively eliminates specific loci by integrating CRISPR-Cas9 mediated Double-Strand Break (DSB) generation and homology-directed recombination with yeast sexual assortment. The method enables **M**arker-less **E**nrichment and **R**ecombination of **G**enetically **E**ngineered loci (MERGE) in yeast. We show that MERGE converts single heterologous yeast loci to homozygous loci at ∼100% efficiency, independent of chromosomal location. Furthermore, MERGE is equally efficient at converting and combining loci, thus identifying viable intermediate genotypes. Finally, we establish the feasibility of MERGE by engineering a fungal carotenoid biosynthesis pathway and most of the human α proteasome core into yeast. MERGE, therefore, lays the foundation for marker-less, highly efficient, and scalable combinatorial genome editing in yeast.

## Introduction

Baker’s yeast has long served as a convenient chassis for bioengineering owing to its genetic tractability, versatile metabolism, and ease of culture in the lab. Decades of fundamental research, together with the development of high throughput toolkits and genome engineering capacities, have established yeast as an ideal model eukaryote for system genetics and synthetic biology^1^. The availability of many selectable genetic markers and simple conversion between haploid and diploid forms has provided avenues to easily combine pairs of genetically engineered loci to understand gene-gene interactions at a global scale^2^. For more extensive genetic alterations, yeast’s highly efficient Homologous Recombination (HR) pathway even enables the synthesis of entire chromosomes, although this approach requires iterative use of selection markers and tedious repetitive procedures^3^. Nonetheless, the ability to alter large contiguous segments of genomic loci has many applications, such as genome minimization^4^, multiplex genome editing^5^, and the total synthesis of the *Mycoplasma* and *E. coli* genomes using yeast HR^6, 7^.

In spite of the progress in whole genome engineering, editing intermediate numbers of independent genomic loci–greater than two and in discontiguous regions of the genome–still presents a significant challenge. While strategies exist for *E. coli*^8^ (e.g., MAGE), diploid organisms such as yeast present the additional editing challenge of multiple alleles for each (independently assorting) genomic locus. Moreover, while high throughput cloning strategies and the reduced cost of *de novo* DNA synthesis now allow swapping entire heterologous pathways or protein complexes into yeast, such efforts frequently require the deletion of corresponding yeast loci, such as the case for efforts to systematically humanize yeast genes^9, 10^, which may entail replacing genes at their corresponding genomic loci to retain native regulation. Beyond these aspects, the expression of each new gene often reveals incompatibilities associated with the engineered pathway. Thus, there is a need for rapid, multi-site, progressive genome editing strategies to address these issues.

The efficiency and speed of CRISPR-Cas9-based genome engineering allow straightforward editing of multiple yeast loci in a single strain, eliminating the need for markers^11–15^. However, the approach gets progressively more challenging for multi-gene systems when the fitness of the intermediate genotypes is unknown (**Figure S1**). Therefore, rationally building heterologous genetic modules in a yeast surrogate requires a highly scalable and combinatorial genome editing technology.

Synthetic Genetic Array (SGA) analysis permits the combination of loci in a fitness-driven manner, requiring markers linked to the modified loci and haploid-specific selections^16^. Using an SGA-like strategy to build a heterologous multi-gene system would need several unique markers linked to each gene. However, the lack of adequate selection markers limits its application. The Green Monster (GM) method bypasses the marker dependency by using Green Fluorescent Protein (GFP) expression as a readout to combine engineered loci^17^. However, implementing the GM strategy can be challenging for heterologous pathway engineering with an added burden of expressing fluorescent cassettes^18^. Thus, accomplishing large-scale combinatorial genome editing in yeast necessitates new genetic tools that circumvent these requirements.

In this work, we describe a novel, CRISPR-Cas9-based method to readily combine genetically engineered loci without the need for markers and exogenous repair templates. The approach involves generating CRISPR-Cas9-mediated Double-Strand Breaks (DSBs) and highly efficient Homology Directed Repair (HDR) along with yeast mating and sporulation to randomly assort edited sites performing multi-site and marker-less combinations of engineered loci. This selective and successive elimination of specific yeast loci mimics a gene drive^19–22^ (**Figure S2**) and facilitates Marker-less Enrichment and Recombination of Genetically Engineered yeast loci (**MERGE**).

We show that CRISPR-mediated selection operates similarly to classical selection markers, enabling MERGE to efficiently explore many combinations of genetically engineered loci, revealing a fitness-driven path to engineering any heterologous system in yeast. We further demonstrate that MERGE enables rapid assembly of an entire carotenoid biosynthesis pathway by performing a multi-site and marker-less combination of distinct engineered loci. Finally, using a multiplexed version of MERGE, we humanize a near-complete α proteasome core (6 of 7 subunits) in yeast while revealing a fitness-driven path to the humanization of complex processes.

## Results & Discussion

### CRISPR-Cas9 allows marker-less selection and enrichment of unique genotypes in yeast (CELECT)

CRISPR-Cas9 enables the precise and marker-less editing of both essential and non-essential yeast loci owing to significantly lower error-prone Non-Homologous End Joining (NHEJ) in yeast relative to HDR^23^. Therefore, Cas9-sgRNA-induced lethality serves as a rapid test for a functional CRISPR reagent in yeast (Colony Forming Units Observed or **CFU_O_** = ∼0) compared to the Cas9 alone (Colony Forming Units Expected or **CFU_E_**). To verify the ON-target activity of CRISPR reagents, the transformation of a plasmid carrying expression cassettes for both Cas9 and a sgRNA targeting a specific locus (**pCas9-sgRNA*^locus^***) in a yeast strain that harbors a corresponding engineered locus resistant to further targeting allows the survival of colonies similar to the pCas9 alone (**CFU_O_**/**CFU_E_** = **∼1**) (**Figure 1A**).

**Figure 1.**
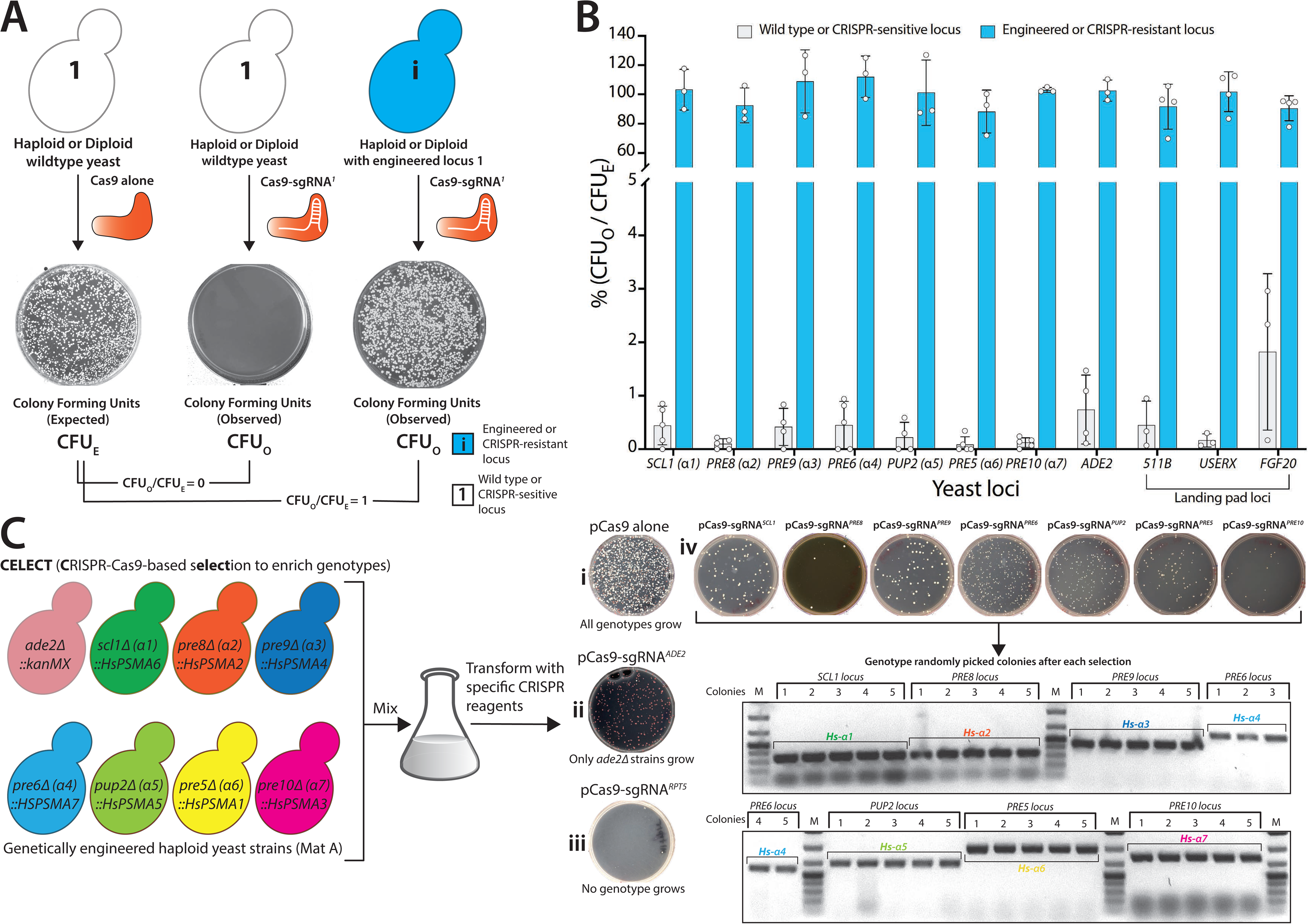
CRISPR-mediated selection (CELECT) enables the enrichment of unique genotypes. **(A) pCas9-sgRNA*^1^*** targeted to any yeast locus (locus **1**) leads to lethality (**CFU_O_**) compared to the vector without sgRNA (pCas9 alone; **CFU_E_**). However, the modification of locus **1** to **i** prevents further targeting by the corresponding CRISPR reagent (**CFU_O_**). A simple readout of **CFU_O_/CFU_E_** enables the identification of modified genotypes. **(B)** Each CRISPR reagent targeted to several wild-type yeast loci shows near 100% lethality (Gray bars or CRISPR-sensitive loci). However, after editing the loci (by generating single-humanized yeast α proteasome genes, *ade2Δ::kanMX* and inserting carotenoid genes at landing pad loci), the strains show resistance to the corresponding CRISPR reagent (Blue bars or CRISPR-resistant loci, **CFU_O_/CFU_E_ = ∼1**). **(C)** Each unique genetically modified strain can be enriched from the mixture using the corresponding CRISPR reagent (CELECT). The transformation of pCas9 alone serves as a control (**i**). Haploid *ade2Δ::kanMX* strain quantifies the efficiency of the CRISPR selection (**pCas9-sgRNA*^ADE2^***), enriching only the *ade2Δ* homozygous genotypes (red-colony phenotype) (**ii**). The transformation of **pCas9-sgRNA*^RPT5^*** causes lethality as all the strains in the mix harbor a wild-type copy of the *RPT5* gene (**iii**). Similarly, each CRISPR reagent specific to individual yeast *α* proteasome genes also selected the corresponding humanized strains as demonstrated by PCR-based genotyping of randomly picked colonies (**iv**).

We tested this strategy by humanizing α proteasome core genes in yeast that are functionally replaceable by their human counterparts^24^. Using CRISPR-Cas9, we replaced each yeast α proteasome gene with its human ortholog at the native loci (**Figure S3**). Additionally, we modified non-essential loci to test if the CRISPR-mediated selection is broadly applicable and found that all engineered strains are resistant to the corresponding CRISPR-Cas9-sgRNA mediated lethality (**CFU_O_**/**CFU_E_ = ∼1, Figure 1B, Figure S4A & S4B**). Henceforth, we refer to the <u>C</u>RISPR-Cas9-mediated s<u>elect</u>ion of genotypes as CELECT.

To further verify if the resistance of humanized strains is exclusive to its corresponding CRISPR reagent, all single-humanized α proteasome strains were mixed in culture (**Figure 1C**). To quantify the enrichment of unique genotypes, we inoculated a *ade2Δ::kanMX* haploid strain in the mixture. The **pCas9-sgRNA*^ADE2^*** exclusively enriched for resistant *ade2Δ* genotype, whereas all other genotypes in the mix harboring a wild type *ADE2* locus are inviable (**Figure 1C-ii**). Conversely, the transformation of **pCas9-sgRNA*^RPT5^*** targeting *RPT5* (a base subunit of the proteasome complex), for which all the strains in the mix harbor a wild-type copy, shows no survivors (**Figure 1C-iii**). We demonstrate that each CRISPR reagent targeting yeast α proteasome genes selected a corresponding humanized genotype from a mix, respectively (**Figure 1C-iv**). Thus, CRISPR-Cas9-mediated resistance functions similarly to conventional antibiotic or auxotrophic markers in yeast.

### MERGE^0^ is nearly 100% efficient at converting loci irrespective of the yeast gene location on the chromosome

Mating of engineered haploid strains with wild-type generates a heterozygous genotype. CRISPR-Cas9 reagent targeted to either allele should enable the conversion to homozygous diploid at high efficiency while also enriching the desired genotype (**MERGE level 0** or **MERGE^0^**) (**Figure 2A**). We used *ADE2/ade2Δ::kanMX* heterozygous knockout diploid (hetKO) strain to quantify the efficiency of **MERGE^0^**. The wild type *ADE2* allele is susceptible to **pCas9-sgRNA*^ADE2^*** mediated DSB. In contrast, the *ade2Δ::kanMX* allele is resistant, proving a readout of conversion as the loss-of-function of *ADE2* results in a red color colony phenotype.

**Figure 2.**
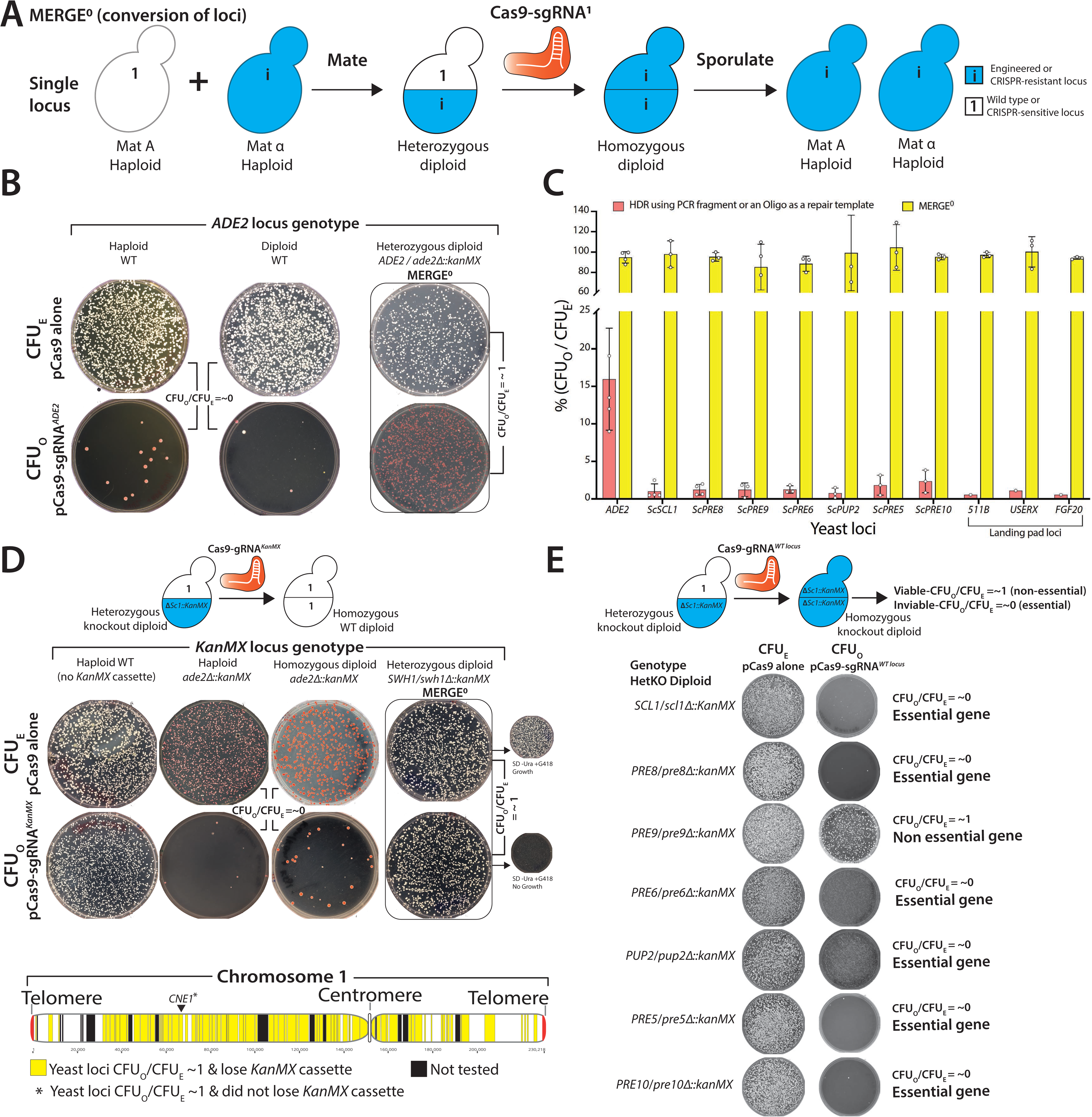
MERGE level 0 (MERGE^0^) efficiently converts a single heterozygous to a homozygous locus. **(A)** Mating haploid yeast strains, each harboring a different allele at a single locus, enables the combination as a heterozygote. CRISPR reagent targeted to one of the alleles (CRISPR-sensitive locus) initiates recombination using the homologous chromosome with a CRISPR-resistant locus as a repair template. Sporulation of the resulting homozygous diploid yields the desired genotype in both mating types. **(B)** CRISPR reagent (**pCas9-sgRNA*^ADE2^***) targeted to the wild type *ADE2* locus quantifies the efficiency of **MERGE^0^**. The transformation of **pCas9-sgRNA*^ADE2^*** in the wild type haploid or diploid strains is lethal, with few surviving red colonies suggesting efficient ON-target activity (**CFU_O_/CFU_E_ = ∼0**). However, the transformation of **pCas9-sgRNA*^ADE2^*** in heterozygous diploid *ADE2/ade2Δ::kanMX* strain shows no lethality (**CFU_O_/CFU_E_ = ∼1**). Instead, all surviving red colonies suggest efficient conversion to the knockout locus. **(C)** Similarly, CRISPR reagents targeted to several wild-type yeast loci show near 100% lethality with few surviving colonies while simultaneously providing exogenous repair templates (oligo for *ADE2* locus and PCR fragments for the remaining loci) (Red bars). However, each heterozygous diploid displays resistance to CRISPR reagents respectively (**CFU_O_/CFU_E_ = ∼1,** Yellow bars). PCR confirmation verified the conversion to the CRISPR-resistant locus. **(D)** CRISPR reagent targeting *KanMX* cassette (**pCas9-sgRNA*^KanMX^***) shows no OFF-target activity in a strain lacking the cassette (**CFU_O_/CFU_E_ = ∼1**). However, haploid or diploid strains harboring *KanMX* cassettes as the only allele show lethality (**CFU_O_/CFU_E_ = ∼0**), suggesting ON-target activity. However, diploid strain heterozygous for *KanMX* allele (*SWH1*/*swh1Δ::kanMX*) is viable (**CFU_O_/CFU_E_ = ∼1**), suggesting conversion to the wild type allele as demonstrated by the loss of G418 resistance after performing **MERGE^0^**. Using heterozygous diploid knockout strains (hetKO) arrayed across the entire yeast **chromosome I**, **MERGE^0^** similarly converted every knockout allele to the wild type locus (Yellow = **CFU_O_/CFU_E_ = ∼1**), except in the case of *CNE1* that showed resistance to the CRISPR reagent but did not lose the *KanMX* cassette. **(E)** Alternatively, using hetKO strains and targeting CRISPR reagent to the wild-type yeast locus enables a single-step gene essentiality assay in yeast. The remarkably high efficiency of **MERGE^0^** converts every heterozygous yeast locus to homozygous null, therefore, allowing viability only if the gene is non-essential (as in the case of yeast proteasome *α*3 gene). All other essential loci show lethality after **MERGE^0^**.

The transformation of the wild-type haploid or diploid yeast with **pCas9-sgRNA*^ADE2^*** showed a lethal phenotype (∼0-20 **CFU_O_** on average) (**Figure 2B**). However, **pCas9-sgRNA*^ADE2^*** transformation in *ADE2*/*ade2Δ*::*kanMX* hetKO strain shows resistance while also converting the locus to *Δade2*::*KanMX* at ∼100% efficiency (**Figures 2B & S5A**). **MERGE^0^** performs with a comparable proficiency at single-humanized α proteasome and landing pad loci (**Figures 2C, S5B & S5C**). To further test if **MERGE^0^** can convert any heterozygous yeast locus independent of the position on a chromosome, we explored the strategy across many yeast genes located on chromosome 1. We designed a CRISPR reagent to target *KanMX* (**pCAS9-sgRNA*^KanMX^***) in hetKO diploid strains harboring the *KanMX* cassette instead of a yeast gene. **MERGE^0^** converted all heterozygous loci to homozygous wild-type alleles, respectively, with simultaneous loss of *KanMX* cassette (except for *CNE1*) (**Figure 2D, S6A & S6B**).

Furthermore, using CRISPR-Cas9 to target a wild-type yeast locus instead of the *KanMX* cassette in a hetKO diploid strains, **MERGE^0^** enables single-step gene essentiality screening in yeast, significantly increasing efficiency compared to the classical methods such as Tetrad dissection or SGA analysis^25, 26^. Except for a non-essential *α3* gene, all six essential α proteasome genes are inviable post-transformation of the corresponding CRISPR reagents (**Figure 2E**).

### MERGE^1^ permits a fitness-driven combination of the engineered loci

Given the high efficiency of **MERGE^0^** at converting one yeast locus, we next tested if the method works similarly to convert and combine two distinct loci (**MERGE level 1 or MERGE^1^**). Mating yeast strains, each with one modified locus, facilitate the combination as heterozygotes at two separate loci. CRISPR-Cas9 with two sgRNAs that target each unique locus allows simultaneous conversion to paired CRISPR-resistant loci, respectively (**Figure 3A**). To test the pipeline across all humanized *α* proteasome strains, we first used **MERGE^0^** to move all singly humanized *α* proteasome loci from BY4741 background to SGA strain background in both mating-types (**Figure S4B**). To verify if **MERGE^1^** can simultaneously convert two distinct loci, we tested the efficiency of combining paired-humanized α3/α7 and α1/α7 genotypes. We show that **MERGE^1^** is highly efficient as each heterozygous genotype is resistant to the double-sgRNA CRISPR-Cas9 reagent while converting and combining two humanized loci (**Figure 3B, S7A**). The CRISPR reagent is lethal in wild-type and singly humanized strains (**Figure 3B & S7B**). The sporulation of viable diploids yields haploid double-humanized strains in both mating-types. Genotyping randomly picked colonies confirms the conversion to humanized loci (**Figure 3B**). Additionally, the double-humanized α7/α1 genotype, while viable, shows a sporulation defect (a phenotype associated with *Hsα7*) (**Figure S7A**). Thus, **MERGE^1^** allows the survival and enrichment of only viable homozygous double-humanized loci without the requirement of any diploid-specific selection. However, a double-humanized α5/α7 is inviable as a combined genotype, as evidenced by double-sgRNA CRISPR-Cas9-mediated lethality (**CFU_O_/CFU_E_ = ∼0**). PCR-based genotyping of a surviving colony shows homozygosity of only the Hsα7 locus (**Figure 3C**). In contrast, while a single-humanized Hs*α6* strain is temperature-sensitive (TS) at 37°C, combination with the neighboring humanized α7 gene rescues the phenotype (**Figure S7A′**). However, the TS phenotype is associated with one of the variants of Hsα6 (Variant 1, 37>Glycine) (**Figure S8A**), whereas another common variant (Variant 2, 37>Valine) shows no growth defect at 37°C (**Figure S8B**). Thus, **MERGE^1^** revealed the fitness of paired-humanized genotypes similar to synthetic genetic interactions without the need for linked markers or haploid-specific selections^16^.

**Figure 3.**
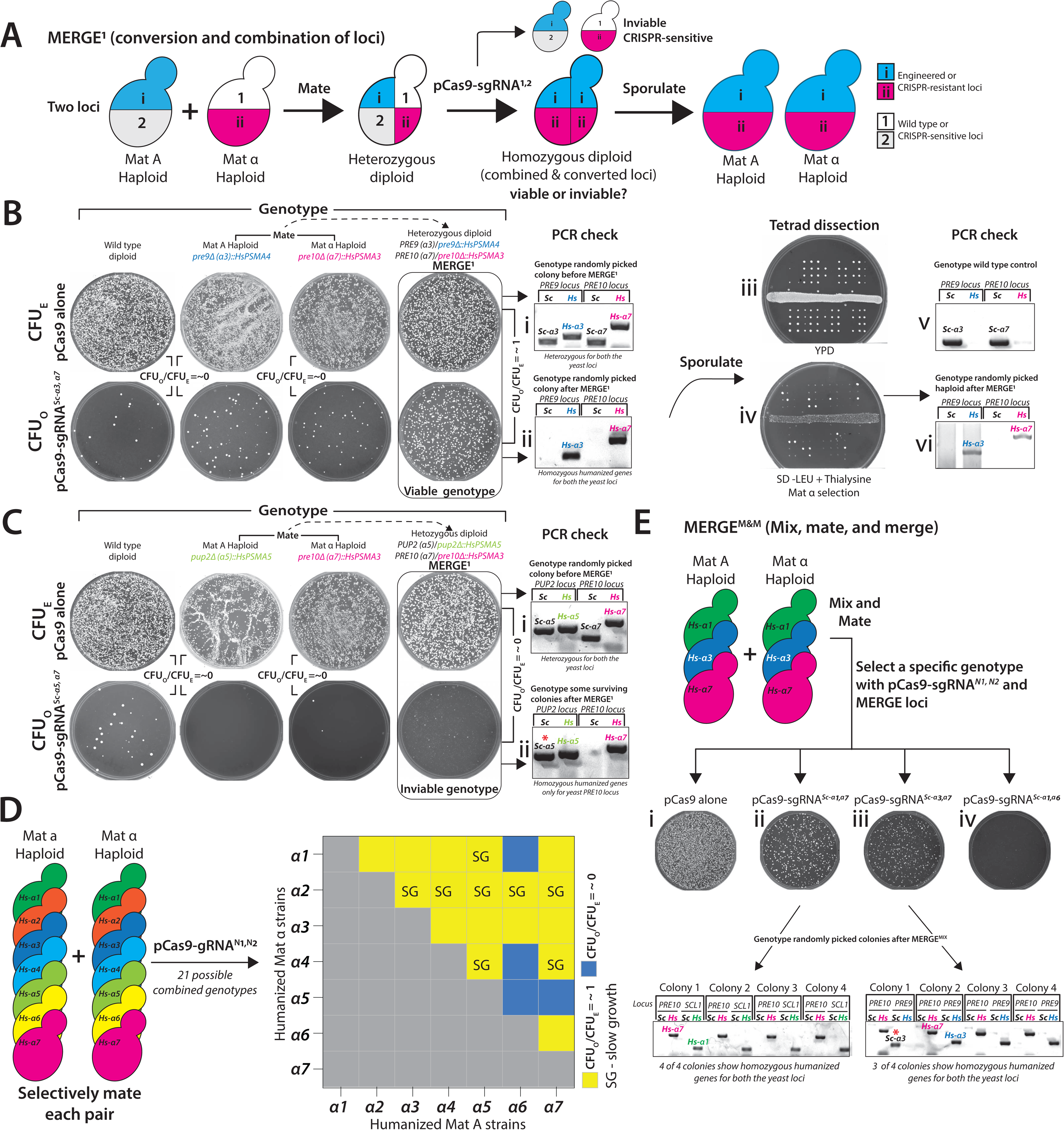
MERGE level 1 (MERGE^1^) efficiently converts and combines two heterozygous to homozygous loci. **(A)** The schematic shows the mating of haploid yeast strains, each harboring two different alleles, enabling the combination as a heterozygote for each locus. Double-sgRNA CRISPR reagent targeted to both the alleles (CRISPR-sensitive loci, **1 & 2**) initiates recombination at both loci using the homologous chromosome with CRISPR-resistant loci (**i & ii**) as a repair template, thus, enabling the simultaneous combination of 2 loci. **CFU_O_/CFU_E_ = ∼1** suggests viable combined genotypes. Sporulation of the resulting homozygous diploid yields the desired combined genotype in both mating types. **(B)** Double-sgRNA CRISPR reagent (**pCas9-sgRNA*^Scα3,α7^***) targeted to two wild type yeast loci (proteasome *α*3 and *α*7 genes) quantifies the efficiency of **MERGE^1^**. The transformation of **pCas9-sgRNA*^Scα3,α7^*** in the wild type, and single-humanized haploid Hs*α*3 or Hs*α*7 strains is lethal, with few surviving colonies (**CFU_O_/CFU_E_ = ∼0**). However, the transformation of **pCas9-sgRNA*^Scα3,α7^*** in a heterozygous diploid humanized strain shows no lethality (**CFU_O_/CFU_E_ = ∼1**). PCR-based genotyping of surviving diploid strains after **MERGE^1^** shows conversion of both yeast to the humanized loci (**ii**) compared to before **MERGE^1^** (**i**). Sporulation generated the paired-humanized haploids in both mating-types (shown here as Tetrads spotted on YPD (**iii**) and Mat *α* selection-**iv**). PCR check of haploid strains confirms the combination of humanized loci (**vi**). PCR of wild-type yeast loci is shown as a reference (**v**). **(C)** Alternatively, double-sgRNA CRISPR reagent (**pCas9-sgRNA*^Scα5,α7^***) targeting two wild type yeast loci (proteasome *α*5 and *α*7 genes) in a corresponding heterozygous diploid humanized strain showed lethality, suggesting incompatible combination (**CFU_O_/CFU_E_ = ∼0**). The transformation of **pCas9-sgRNA*^Scα5,α7^*** in the wild type, and single-humanized Hs*α5* or Hs*α7* strains is lethal, with few surviving colonies, suggesting ON-target activity (**CFU_O_/CFU_E_ = ∼0**). PCR-based genotyping of surviving colonies after **MERGE^1^** shows the conversion of only one (*α5*) yeast to the humanized locus (**ii**) compared to before **MERGE^1^** showing heterozygous genotype at both loci (**i**). **(D) MERGE^0^** generated 7 humanized *α* proteasome strains in each mating-type. Mating each single-humanized strain allows a systematic test for every viable double-humanized genotype (21 combined genotypes). Transformation of double-sgRNA CRISPR reagent uniquely targeting yeast genes in each paired heterozygote facilitates the combination of two humanized loci as homozygotes (**CFU_O_/CFU_E_ = ∼1, yellow**) while also revealing genotypes that are not permitted (**CFU_O_/CFU_E_ = ∼0, blue**). **(E)** Mixing and mating singly humanized genotypes (Hs*α*1, Hs*α*3 and Hs*α*7) allows a random combination of two humanized alleles after double-sgRNA CRISPR selection referred to as mix, mate, and MERGE (**MERGE^M&M^**). Transformation of **pCas9 alone** serves as a control (**i**). Double-sgRNA CRISPR reagents, **pCas9-sgRNA*^Scα1,α7^*** (**ii**) and **pCas9-sgRNA*^Scα3,α7^*** (**iii**), specifically enriched the corresponding double-humanized genotype while also converting yeast to humanized loci. PCR-based genotyping of randomly picked colonies confirms *Hsα1α7* (4 of 4 colonies tested) and *Hsα3α7* (3 of 4 colonies tested). Comparatively, the transformation of double-sgRNA CRISPR reagent, **pCas9-sgRNA*^Scα1,α6^*** (**iv**), shows no surviving colonies as the genotype does not exist in the mixture.

To systematically determine if there are specific pairwise restrictions to the humanization of α-subunits, we mated all haploid single-humanized strains obtaining diploid heterozygous genotypes (21 different genotypes). The corresponding CRISPR-Cas9-based selection and a simple readout of **CFU_O_/CFU_E_** identified the permitted double-humanized genotypes (17/21 genotypes). The data reveals that only specific double-humanized genotypes are viable, whereas some are not (**Figure 3D & S7C**). The incompatibility of paired genotypes comprising *Hsα1/α6*, *Hsα4/α6*, *Hsα5/α6*, and *Hsα5/α7* may likely be due to the missing neighboring interactions within the α proteasome core, except in the case of *Hsα5,α6* pair (**Figure S7C′**). A sequential editing strategy successfully engineered some paired-humanized genotypes to confirm that **MERGE^1^** represents viable/fit paired genotypes. However, it did not provide a clear perspective of incompatibilities (**Figure S7D**).

Given the success of **MERGE^1^**, we examined the strategy to select double-humanized genotypes randomly from a mixture (Mix, Mate & MERGE or **MERGE^M&M^**) (**Figure 3E**). We inoculated single-humanized *α1, α3* and *α7* haploid strains of both mating-types as a mix, allowing random mating. Each double-sgRNA CRISPR-Cas9 selection enriched the corresponding paired genotype from the mated mix while converting the wild-type yeast to humanized loci (**Figure 3E-iii & 3E-iv**). In mixed culture, the transformation of **pCas9-sgRNA*^Sc-α1,α6^*,** a selection for a non-existing genotype, does not yield any viable genotype (**Figure 3F-ii**). Therefore, **MERGE^M&M^** can be scaled to obtain several paired genotype combinations of engineered loci from a mix.

### MERGE is scalable to combine multiple genetically engineered loci

The CRISPR-Cas9 system enables multiplexed editing by introducing multiple sgRNAs^27^. To test scalability, we designed **MERGE^MX^** (**MERGE, mate and multiplex**) to verify if >2 genetic loci can simultaneously convert to engineered loci by building the 4-gene carotenoid biosynthesis pathway from the carotenogenic yeast *Xanthophyllomyces dendrorhous* into Baker’s yeast (**Figure 4A**). The carotenoid pathway provides a colony color readout as a proxy for pathway engineering (**Figure 4A**). **MERGE^0^** generated haploid strains of opposite mating-types for each carotenoid transcription unit (**Figure S5C**), **MERGE^1^** provided the double-carotenoid genotypes (**Figure 4B**), and **MERGE^MX^** enriched a complete homozygous carotenoid pathway genotype (**Figure 4C**). Furthermore, genotyping of randomly picked dark orange colonies confirmed the conversion and combination of the engineered loci (**Figure 4C**).

**Figure 4.**
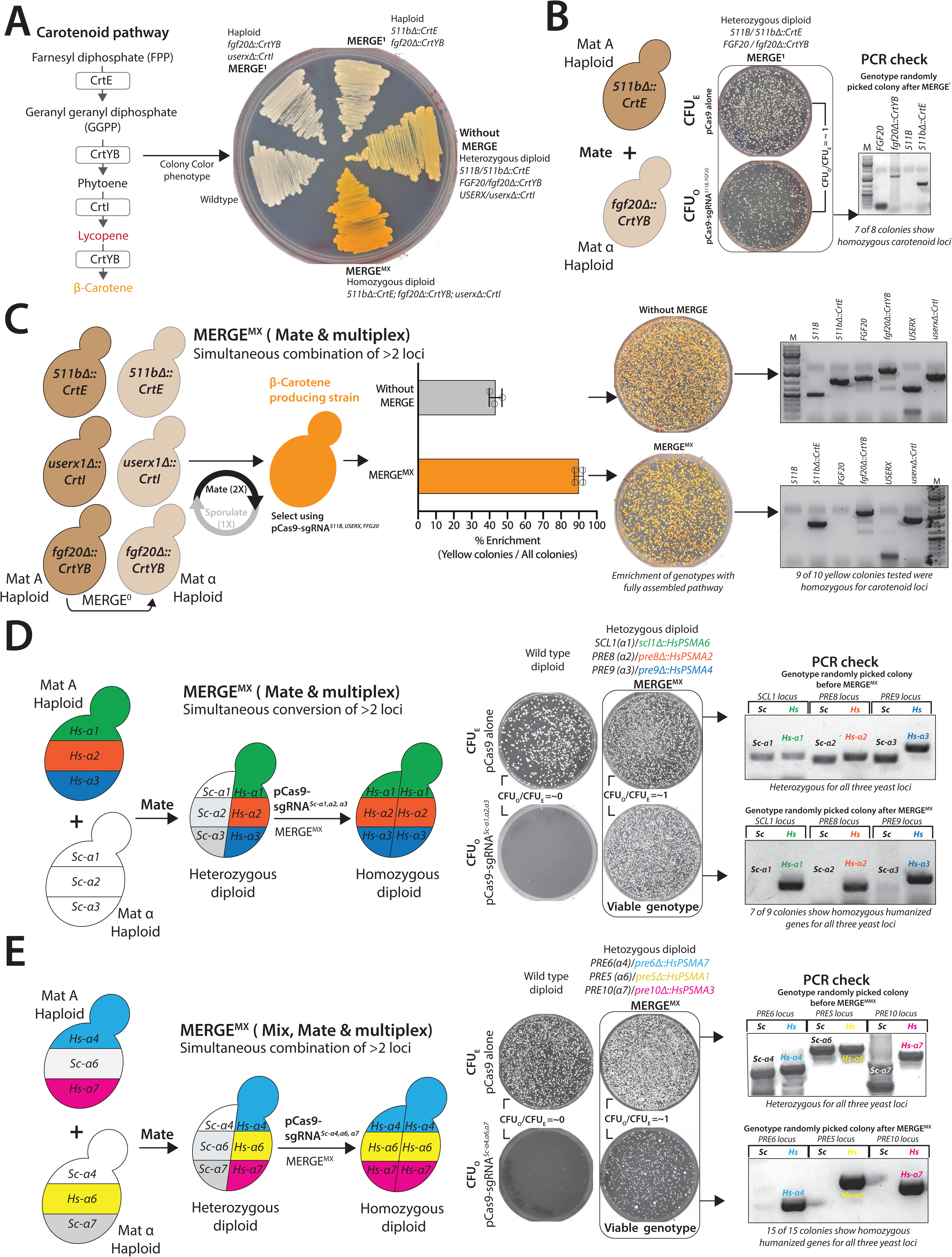
MERGE mate and multiplex (MERGE^MX^) is scalable to combine multiple engineered loci. **(A)** Schematic shows the carotenoid pathway genes and the metabolic intermediates leading to the color colony phenotype. A complete pathway (*CrtE*, *CrtYB* & *CrtI*) leads to an orange colony appearance, whereas the partial assembly (*CrtYB* & *CrtI*) provides an off-white colony phenotype. **MERGE^1^** provided single-carotenoid engineered strains in both mating types. The homozygous diploid for a complete carotenoid pathway shows an intense orange colony phenotype than the heterozygous strain. **(B)** Schematic showing **MERGE1** combining two engineered genotypes at landing pad loci. Double-sgRNA CRISPR reagent (**pCas9-sgRNA*^511B,FGF20^***) targeting two landing pad loci (*511B* and *FGF20*) quantifies the efficiency of **MERGE^1^**. The transformation of **pCas9-sgRNA*^511B,FGF20^*** in double heterozygous diploid strain shows no lethality (**CFU_O_/CFU_E_ = ∼1**). PCR- based genotyping of several colonies after **MERGE^1^** shows the conversion to the engineered carotenoid loci. **(C)** Single-carotenoid gene strains of both mating-types were mixed, mated (2X) and sporulated (1X). The mixture transformed with **pCas9 alone** shows an equal distribution of white, off-white and orange-colored colonies. However, the triple-sgRNA **pCas9-sgRNA*^511B,FGF20,USERX^*** transformed mix significantly enriched the yellow colonies. PCR-based genotyping of several colonies showed **MERGE^MX^** successfully homozygosed all loci, whereas the yellow colonies on the unselected plate showed heterozygous loci. **(D)** Schematic shows mating haploid triple-humanized Hs*α*1,*α*2,*α*3 strain with a wild-type yeast enables the combination of 3 humanized loci as heterozygotes. Triple-sgRNA CRISPR reagent (**pCas9-sgRNA*^Scα1,α2,α3^***) targeted to the corresponding wild-type yeast alleles (CRISPR-sensitive loci) simultaneously initiates recombination at all 3 loci using the homologous chromosome with humanized CRISPR- resistant loci as a repair template. The transformation of **pCas9-sgRNA*^Scα1,α2,α3^*** in the wild type is lethal, suggesting ON-target activity (**CFU_O_/CFU_E_ = ∼0**). However, the transformation of **pCas9-sgRNA*^Scα1,α2,α3^*** in a diploid triple heterozygous humanized strain shows no lethality (**CFU_O_/CFU_E_ = ∼1**). PCR-based genotyping of surviving colonies after **MERGE^MX^** confirms the conversion of all 3 yeast to the humanized loci compared to before **MERGE^MX^**. The transformation of a plasmid without sgRNA (**pCas9 alone**) serves as a transformation efficiency control (**CFU_E_**). **(E) MERGE^MX^** similarly combined >2 loci after mating a strain with 2 humanized loci (Hs*α*4,*α*7, Mat A) with a strain carrying 1 humanized locus (Hs*α*6, Mat α). The transformation of **pCas9-sgRNA*^Scα4,α6,α7^*** in the wild type is lethal, suggesting ON-target activity (**CFU_O_/CFU_E_ = ∼0**). However, the transformation of **pCas9-sgRNA*^Scα4,α6,α7^*** in a mated mix selected a diploid triple-humanized strain (**CFU_O_/CFU_E_ = ∼1**). PCR-based genotyping of several colonies after **MERGE^MX^** shows conversion of all yeast to the humanized loci.

To test whether **MERGE^MX^** can perform the combination of >2 humanized *α* proteasome genotypes, we mated a previously obtained Hs*α1α2α3* strain (**MERGE^1^**) with a wild-type type strain generating a heterozygous diploid for all three loci. The strain provided a platform to test **MERGE^MX^** by simultaneously combining 3 distinct humanized loci. The triple-sgRNA CRISPR reagent (**pCas9-sgRNA^Sc-^*^α1,α2,α3^***) converted all yeast loci to human versions (7 of 9 colonies tested) (**Figure 4D**). Next, using **MERGE^MX^**, we tested if a triple-humanized Hs*α4α6α7* genotype is viable. Transformation of triple-sgRNA, **pCas9-sgRNA^Sc-^*^α4,α6,α7^*** in a mated mix of haploid Hs*α4α7* (Mat A) and Hs*α6* (Mat *α*) strains allowed simultaneous conversion of three wild-type yeast loci to humanized versions without using any diploid specific selection (**Figure 4E & S11A**). Thus, if a triple-humanized intermediate genotype is viable, **MERGE^MX^** can enrich and combine the specific genotype.

To further address throughput, we designed a CRISPR reagent targeting a GFP expression cassette in a GM strain^17^ (16 GFP loci, **Figure S9A)**. The transformation of **pCas9-sgRNA*^GFP^*** resulted in a few survivors that failed to show GFP expression, likely due to mutations at GFP loci due to NHEJ (**Figure S9B & S9C**), suggesting successful targeting of the majority of GFP cassettes.

### MERGE enables fitness-driven engineering of a near entire human α proteasome core in yeast

To explore the proficiency of **MERGE** for testing many combinations of engineered loci, we asked if an entire yeast heptameric *α* proteasome ring is humanizable. As an alternate strategy, we also tested the feasibility of sequential engineering using repetitive CRISPR selections and exogenous human gene repair templates (**Figure 5A**). The co-transformation of a triple-sgRNA CRISPR reagent (**pCas9-sgRNA^Sc-^*^α1,α2,α3^***) targeting yeast *α*1, *α*2 and *α*3 genes and PCR fragments of human gene repair templates was successful in obtaining a triple-humanized strain (**Figure S10A-i**). A similar strategy generated triple-humanized Hs*α1α2α4* and Hs*α1α3α4* strains but failed to obtain a quadruple-humanized Hs*α4α5α6α7* (**Figure S10A-ii**). Thus, yeast genes are replaceable sequentially either alone or as small-scale simultaneous replacements. Using a triple-humanized genotype (Hs*α1α2α3*), the humanization of yeast *α*4 was successful. Next, we explored Hs*α*5, Hs*α*6 and Hs*α*7 humanizations in parallel; however, we obtained only one quadruple-humanized Hs*α1α2α3α4α7* genotype (**Figure 5A, Figure S10B**). The functional replacement of yeast *α*5 or *α*6 was unsuccessful despite repeated attempts. The plasmid-borne expression Hs*α6* in a quintuple-humanized strain (Hs*α1α2α3α4α7*) resulted in a toxic phenotype (no growth), suggesting that further humanizations are incompatible (**Figure 5A**). Overall, while the sequential strategy was partly successful, it failed to reveal if the inability to humanize yeast α core entirely was due to incompatible genotypes or inefficient genome editing, especially as hybrid human-yeast genotypes show growth defects and reduced transformation efficiencies (**Figure S10C**).

**Figure 5.**
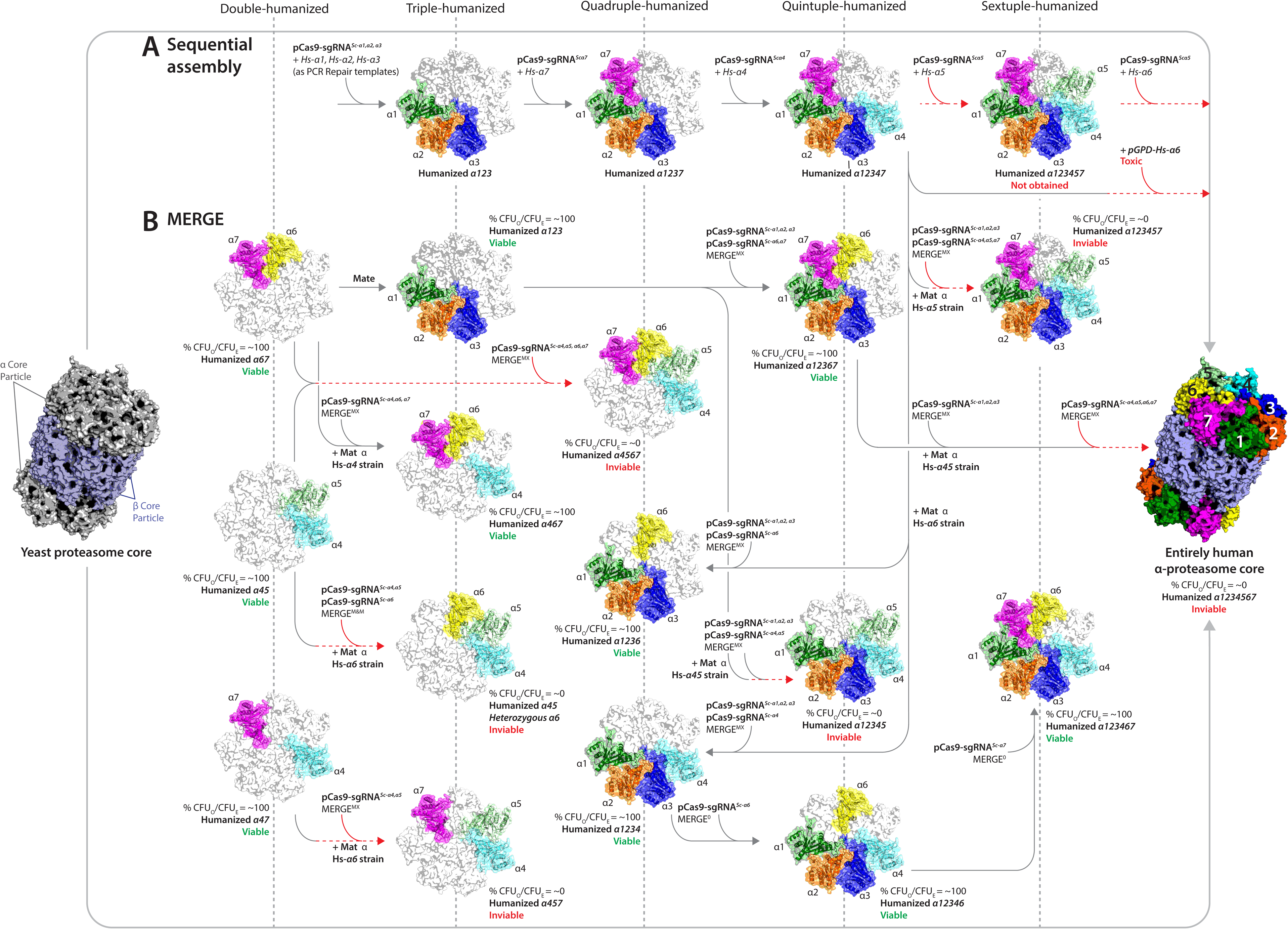
MERGE reveals a fitness-driven path to the humanization of most of the *α* proteasome core in yeast. The schematic shows the transition of heptameric yeast *α* proteasome to humanized *α* proteasome core. **(A)** A sequential strategy used a triple-humanized Hs*α*1,*α*2,*α*3 strain as a background to progressively humanize the rest of yeast *α* core genes by co-transforming a CRISPR reagent and a human gene repair template. Choices to sequentially humanize the *α* proteasome core were made depending on the success of the prior effort. The strategy permitted the humanization of yeast *α*7 followed by α4. Whereas several attempts to humanize yeast *α*5 and *α*6 failed. The expression of human Hs*α*6 in a quintuple-humanized Hs*α*1,*α*2*α*3,*α*4,*α*7 strain resulted in a lethal phenotype. **(B) MERGE** provided a clear readout of the fitness of combined genotypes while revealing incompatible combinations of the humanized yeast α proteasome core. Mating distinct humanized yeast combinations (show as connecting lines) followed by **MERGE^MX^** tested various triple-, quadruple-, quintuple- and sextuple-humanized *α*-proteasome yeast strains. **MERGE** generated viable triple-humanized combinations Hsα4,α5,α6 and Hsα4,α5,α7 (indicated as dashed lines in red), suggesting incompatible combinations. Similarly, while **MERGE^1^** facilitated the generation of Hs*α*4,*α*5 and Hs*α*6,*α*7 genotypes, the subsequent **MERGE^MX^** to generate quadrupled Hs*α*4,*α*5,*α*6,*α*7 failed. Using **MERGE^MX^** followed by **MERGE^0^** identified several quadruples, quintuples and one sextuple-humanized genotype (Hsα1,α2,α3,α4,α6,α7), revealing a fitness-driven path to the humanization of near entire yeast α-proteasome core. Whereas identifying humanized combinations (>2), comprising the Hsα5 subunit, as incompatible genotypes. Mating of Hsα1,α2,α3,α6,α7 and Hsα4,α5 strains generated a heterozygous human-yeast diploid for all 7 α proteasome core genes. Using multiplex-sgRNA CRISPR reagents permitted verifying if the entire human α-proteasome core is feasible. A triple-sgRNA CRISPR reagent (**pCas9-sgRNA*^Sc-α1,α2,α3^***) homozygosed 3 of 7 humanized loci. However, a subsequent **MERGE^MX^,** using a quadruple-sgRNA CRISPR reagent (**pCas9-sgRNA*^Sc-α4,α5,α6,α7^***), failed to obtain a viable genotype. Proteasome core structures were generated using Pymol and PDB- 1RYP. Colored structures show humanized α yeast subunits.

However, **MERGE^MX^** provided a clear readout of incompatible humanized genotypes, readily generating many combinations of humanized genotypes (**Figure 5B**). We first explored several triple-humanized genotypes, obtaining *Hsα*1*α*2*α*3 & *Hsα*4*α*6*α*7 strains (**Figures 4D & 4E**), while *Hsα*4*α*5*α*6 and *Hsα*4*α*5*α*7 genotypes are inviable (**Figures 5B, S11B & S11C**). By mating yeast strains with distinct humanized genotypes, we next explored many higher-order (>3) combinations, obtaining quadruple-humanized Hs*α*1*α*2*α*3*α*4 and Hs*α*1*α*2*α*3*α*7 genotypes. In comparison, the quadruple-humanized *Hsα*4*α*5*α*6*α*7 genotype is not feasible (**Figure 5B, Figure S12A, S12B, S12C & S12D)**. Subsequent **MERGE** strategies generated viable quintuple-humanized genotypes, *Hsα*1*α*2*α*3*α*6*α*7 and *Hsα*1*α*2*α*3*α*4*α*6, whereas *Hsα*1*α*2*α*3*α*4*α*5 genotype is inviable (**Figures 5B**, **S12E, S12F, & S12G)**. The following **MERGE^MX^** assay yielded a viable sextuple-humanized *Hsα*1*α*2*α*3*α*4*α*6*α*7 genotype with a delayed growth phenotype (**Figure S11H**). To conclusively verify if the entire yeast *α* proteasome core is humanizable, we mated partially humanized Hs*α*1*α*2*α*3*α*6*α*7 and Hs*α*4*α*5 strains, allowing the combination of all *α* proteasome genes as heterozygous human-yeast genotypes (**Figure 13A**). **MERGE^MX^** converted yeast *α*1,*α*2, and *α*3 loci to homozygous human alleles. However, the subsequent conversion of the remaining four yeast *α*4*α*5*α*6*α*7 to human loci using **pCas9-sgRNA^Sc-α4,α5,α6,α7^** did not yield viable colonies suggesting a fully human *α* proteasome core is incompatible (**Figure S13B**). Overall, **MERGE** successfully tested many combined humanized genotypes. This gradual progression from yeast to humanized *α* proteasome core rescued the viability of specific incompatible double-humanized genotypes, suggesting that these subunits are co-humanizable when neighboring interactions are restored (**Figures 3D & S14**). The data reveals that the proteasome subunits *α5* and *α6* are not co-humanizable in yeast (**Figure 5**).

Proteasome biogenesis is a highly regulated process aided by several assembly chaperones^28–30^. In the case of *α* proteasome core, particularly *α*5 and *α*6, subunits interact with assembly chaperones, enabling ordered assembly^31–33^. Furthermore, the β core assembly immediately follows the *α* core, and the incompatible interface may now require human β subunits in yeast^28, 34^. The heterologous expression of human assembly chaperones or human β subunits in a humanized *α* core strain may permit the synthesis of a fully human catalytic core particle in yeast. Given the complex assembly and architecture of the proteasome^35^, it is challenging to know if there are a limited number of ‘paths’ to engineer a fully-humanized 20S core particle, due to a rapidly accumulating number of assays to perform (as in **Figure S1**).

The sporulation failure observed in genomically-replaced strains can lead to dead ends while performing MERGE (associated with all humanized Hs*α*7 genotypes, **Figures S11A, S12A & S12F**). We propose two solutions: One, by allowing the strains with heterozygous engineered loci to sporulate without **MERGE**, followed by using **CELECT** to enrich combined haploid genotypes (**Figure S15A**). Alternatively, a sequential strategy can engineer viable genotypes in a haploid strain (**Figures S12F & S15B**). Furthermore, using CRISPR reagents to generate multiple DSBs at several yeast loci could potentially have OFF-target effects. Therefore, we performed whole-genome sequencing of singly humanized (Hs-*α1*) and quintuple-humanized strains (*Hsα*1*α*2*α*3*α4α*7), ruling out OFF-target DSBs and mutations (**Figure S16**).

## Conclusions

In conclusion, MERGE offers highly scalable multi-locus genome engineering in diploid yeast cells by using a high-efficiency CRISPR-based gene-drive-like strategy to overcome the independent assortment of unlinked loci. MERGE has the potential to allow systematic functional genomic analysis in other systems lacking sophisticated tools and drive synthetic and systems biology research from engineering heterologous systems to performing multi-site and genome-wide combinatorial editing in yeast, as we demonstrated by engineering 89 independent sites along Chromosome 1, a complete four gene carotenoid biosynthesis pathway, 16 GFP insertions within the same strain, and a majority of the α proteasome genes. By engineering 6 of 7 human α proteasome core genes in yeast, our work also demonstrates the remarkable degree of functional conservation in the proteasome complex despite over a billion years of evolutionary divergence, extending from a single gene to nearly an entire module. The data confirm our previous observations that humanization seems to be driven by modules of physically or functionally interacting proteins being similarly replaceable^24^. Further characterization of the incompatibilities should reveal novel orthogonal functions or interactions in diverged species. However, pursuing a combinatorial strategy with MERGE along with a sequential strategy in parallel allows one to inform the other about simultaneous replacements that are likely to work. Humanizing all or multiple members of a protein complex will allow a novel approach to learning human biology, including complex assembly, biogenesis, and variant effects on function, investigations of their contributions to disease, and the possibility of seeking therapies for these diseases in the simplified context of a yeast cell.

## Supporting information

Primers used in this study

## Acknowledgements

The authors would like to thank Riddhiman Garge and Dan Boutz (University of Texas at Austin) for many suggestions and critical feedback on gene drives and proteasome biochemistry, Charles Boone (University of Toronto) for sharing the SGA-compatible yeast strains, Frederick Roth (University of Toronto) and Harvard University for sharing Green monster strains, Jef Boeke (New York University) and Grant Brown (University of Toronto) for sharing the carotenoid gene yeast expression plasmids, and Maitreya Dunham (University of Washington) for insightful discussions and feedback on performing MERGE across the entire yeast chromosome.

## Contributions

M.A., B.M.G., J.M.L., A.H.K., and E.M.M. designed the research. M.A., B.M.G., J.M.L., and M.V. performed the experiments. A.H.K. and E.M.M. supervised the research. M.A., B.M.G. A.H.K., and E.M.M. wrote the manuscript with editing help from J.M.L., and M.V.

## Abbreviations

CFU: Colony Forming Units
DSB: Double-Strand Break
HDR: Homology-directed DNA Repair
HR: Homologous Recombination
CELECT: CRISPR-Cas9-based selection to enrich genotypes
MERGE: Marker-less Enrichment and Recombination of Genetically Engineered loci
SGA: Synthetic Genetic Array

---

**Figure S1.**
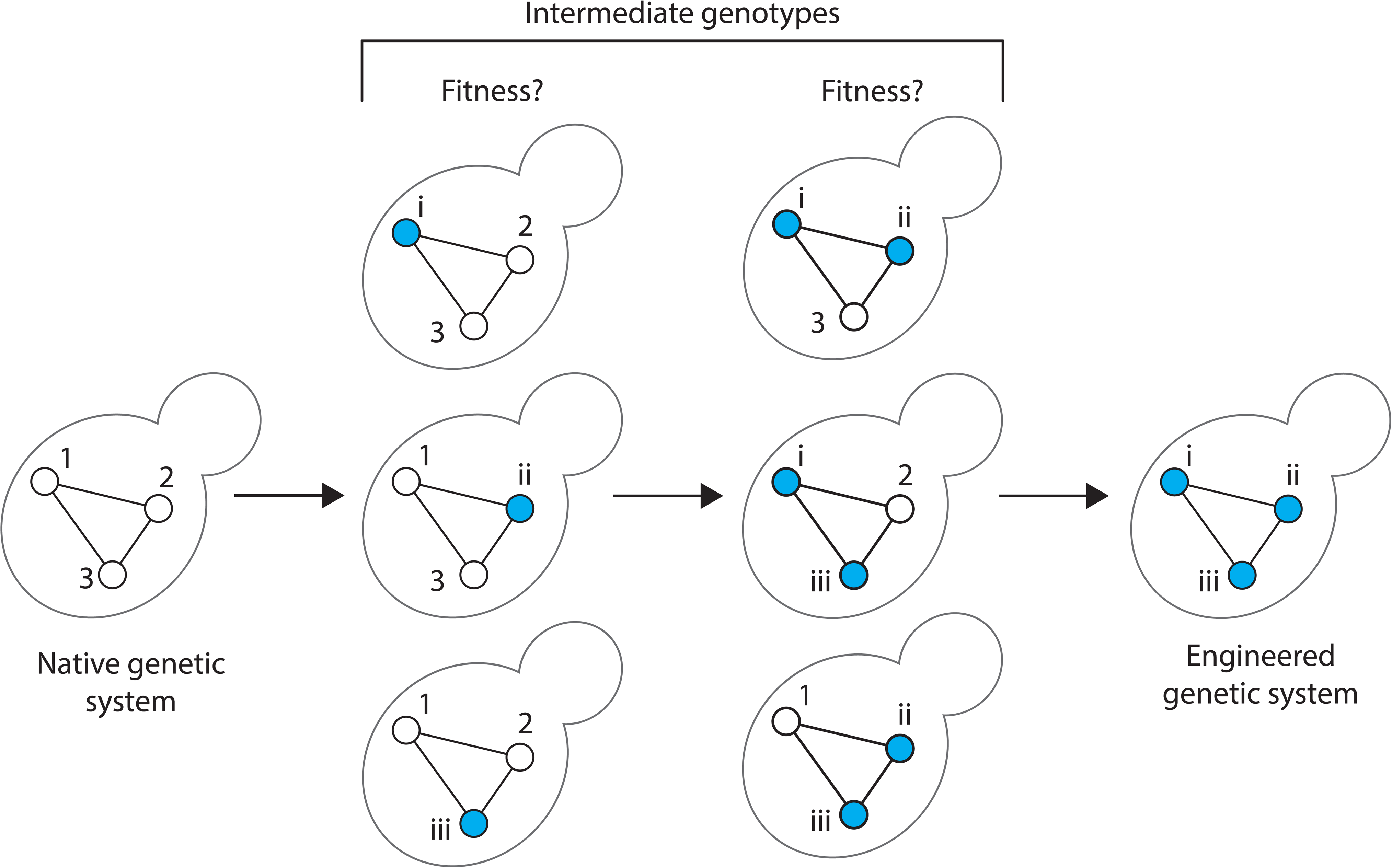
Multiple routes to the engineering of biological systems using a sequential replacement strategy. Let’s examine a simple 3-gene pathway. A sequential replacement strategy to fully engineer this module will navigate 6 possible intermediate genotypes. The following formula can calculate the number of genotypes: *nCr = (n!/n − r!) (r!)*, where n represents a total number of genes to be combined (for example, depicted above, n = 3). And r represents the combined intermediates, like for a single-gene intermediates r = 1 and similarly for a 2-gene intermediates r = 2. Furthermore, while the strategy explores 6 intermediate genotypes, the distinct routes to complete pathway engineering are 9 as some genotypes may require gene 1 and then gene 2, or gene 2 and then gene 1. The higher the number of genes to be engineered, the more the number of intermediate genotypes to be explored. As a reference, for the humanization of the heptameric *α* proteasome core, a total of 126 intermediate hybrid yeast-human genotypes exist towards humanizing the entire yeast *α* core.

**Figure S2.**
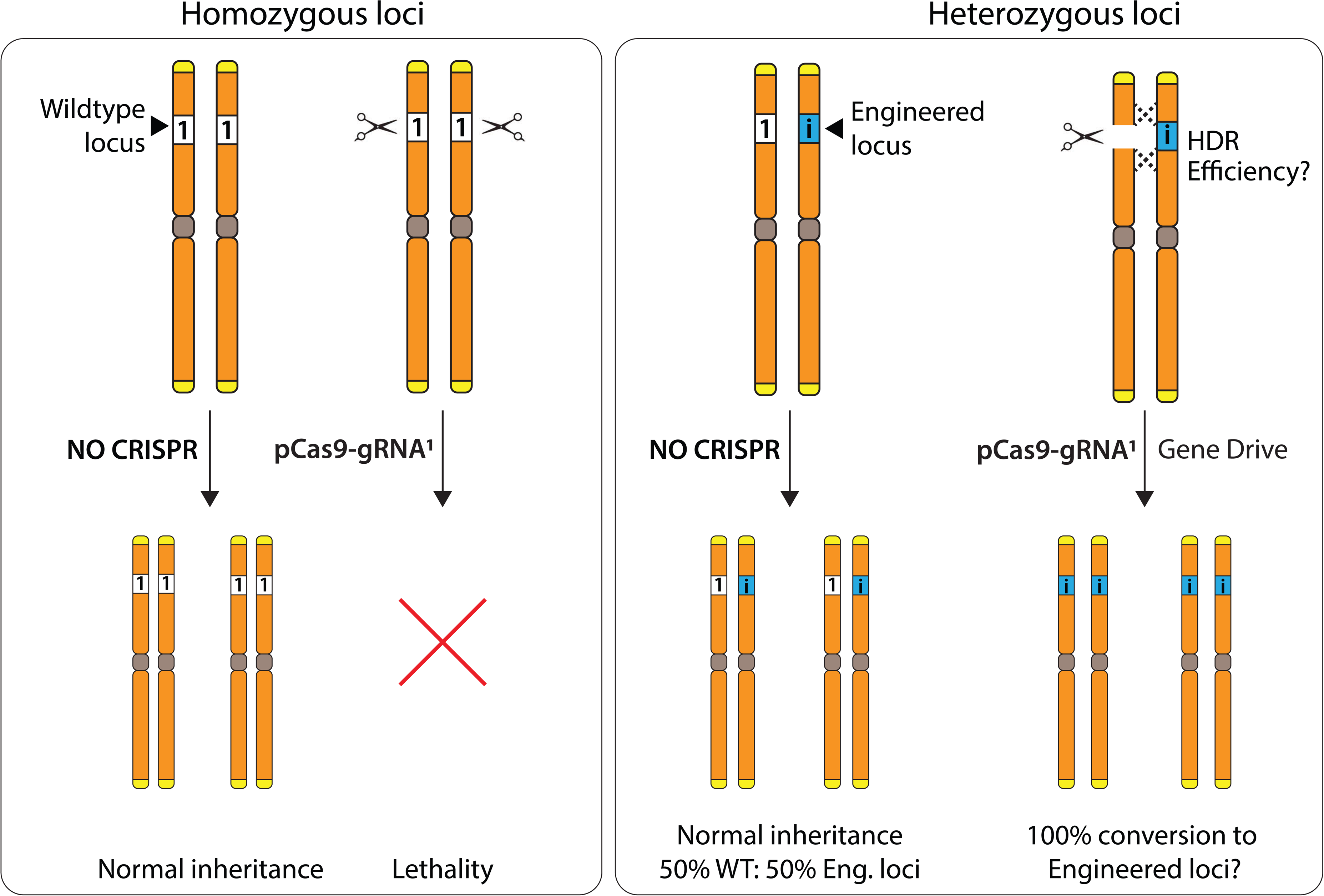
MERGE enables the highly efficient conversion of heterozygous to homozygous loci. Every locus in yeast follows a normal (1:1) inheritance pattern of segregation. However, CRISPR reagent targeted to a haploid or a diploid locus leads to lethality. Similarly, a heterozygous yeast locus follows 1 (1, wild-type) : 1 (i, engineered) inheritance pattern. CRISPR reagent targeted to one of the heterozygous alleles initiates homologous recombination using the CRISPR-resistance homologous chromosome as a repair template, thus enabling efficient conversion to homozygous loci, mimicking a gene drive. The subsequent segregation pattern of 4 : 0 allows propagation of only desired loci.

**Figure S3.**
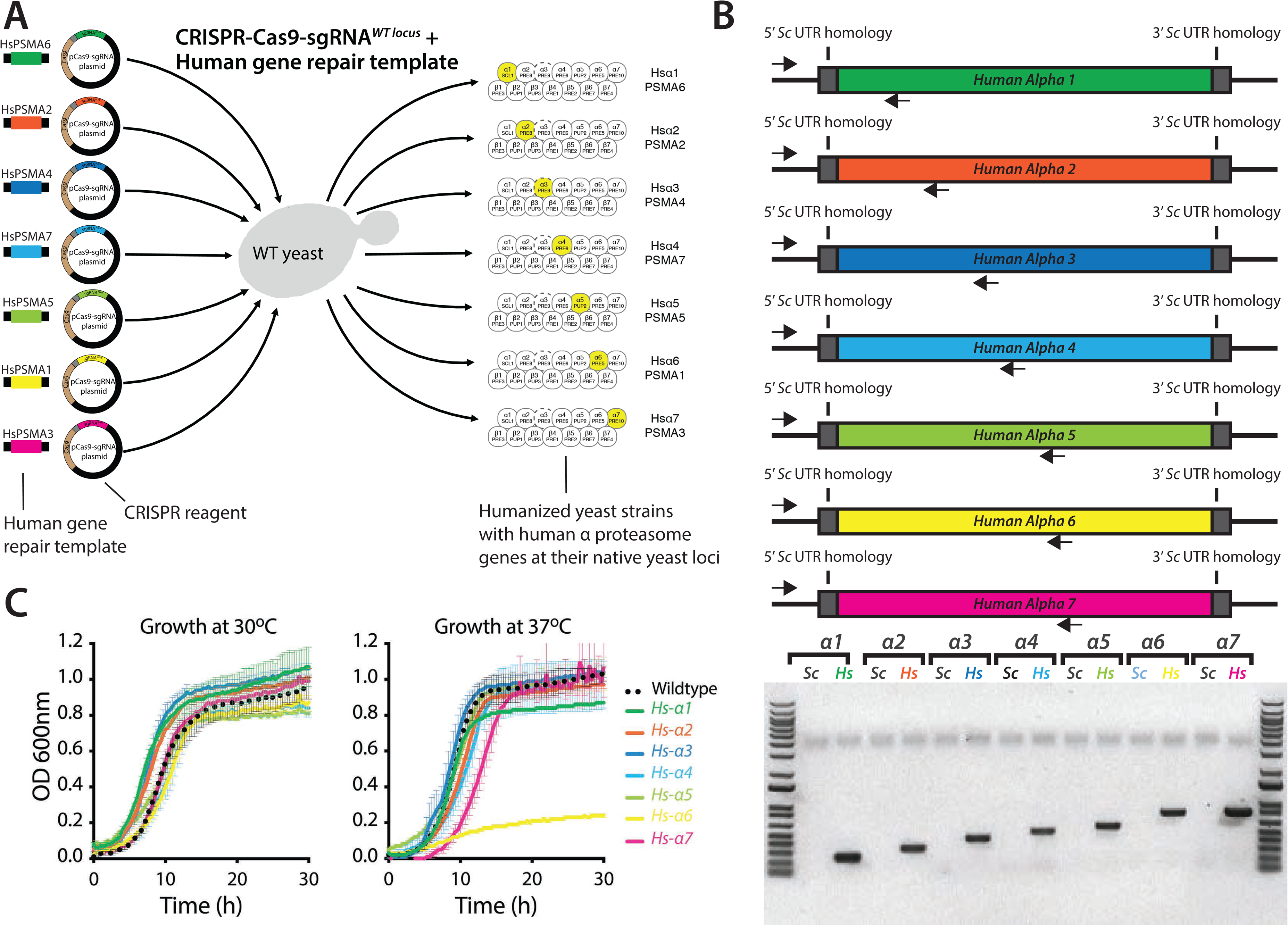
General scheme of CRISPR-Cas9-based gene replacement. **(A)** For each gene, a ‘knockout’ (KO) plasmid (CRISPR reagent) is constructed that expresses the Cas9 endonuclease and a sgRNA with a targeting sequence to the gene of interest. To replace the gene of interest, the CRISPR reagent is co-transformed into yeast with a linear repair DNA, consisting of the replacing gene sequence flanked by long (60-200bp) homology sequences to the flanking region of the genome. Cas9 and the sgRNA are co-expressed and target the gene of interest for cutting, allowing repair by homologous recombination (HR) with the provided repair DNA. **(B)** After obtaining transformants, sequencing across the junction reveals successful replacement (as shown for the human α proteasome subunits replaced at the native yeast loci). PCR for the yeast locus (*Sc*) is negative, whereas for a human gene (*Hs*) is positive. **(C)** The growth of singly-humanized α-subunits is not significantly impaired. Each curve represents an averaging of 4 replicates for each indicated strain, grown in YPD for 24 hours. No strains exhibit significantly impaired growth, though strains with human subunits α4, 5, and 6 show some extension of lag phase at 30°C and marginally slowed doubling times at 30°C. The behavior is similar at 37°C except in *Hs-α6,* which exhibited a significant growth defect.

**Figure S4.**
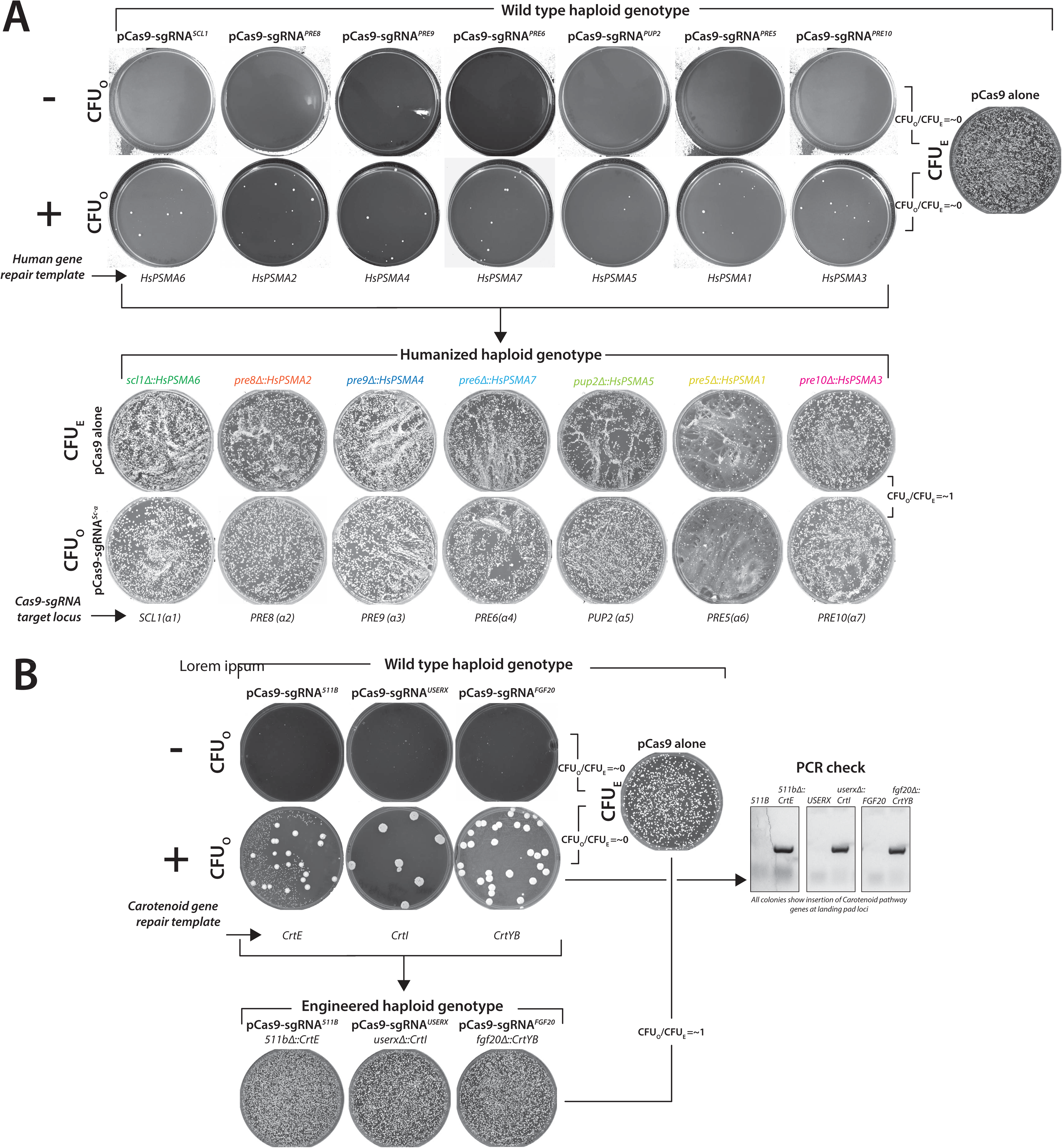
General scheme of CRISPR-Cas9-based gene replacement, CRISPR-mediated sensitivity and resistance of yeast loci. **(A)** CRISPR reagent (**pCas9-sgRNA*^locus^***) targeted to any yeast locus often leads to lethality with few surviving colonies (**CFU_O_**). The corresponding transformation of a control plasmid (**pCas9 alone**) estimates the transformation efficiency (**CFU_E_**). CRISPR reagents targeting 7 yeast α proteasome genes show lethality (**CFU_O_**/**CFU_E_ = ∼0**) with or without the repair template. After humanizing the corresponding yeast loci, each single-humanized haploid strain shows resistance to further targeting by the corresponding CRISPR reagent, respectively (**CFU_O_**/**CFU_E_ = ∼1**). **(B)** CRISPR reagents targeted to landing pad loci (*511B*, *USERX* and *FGF20*) enabled editing by introducing carotenoid gene transcription units *CrtE*, *CrtI* and *CrtYB* (provided as PCR fragment repair templates). The engineered strains show resistance to further cutting by the corresponding CRISPR reagents (**CFU_O_**/**CFU_E_ = ∼1**). PCR-based genotyping confirmed the integration of each carotenoid transcription unit for every colony tested.

**Figure S5.**
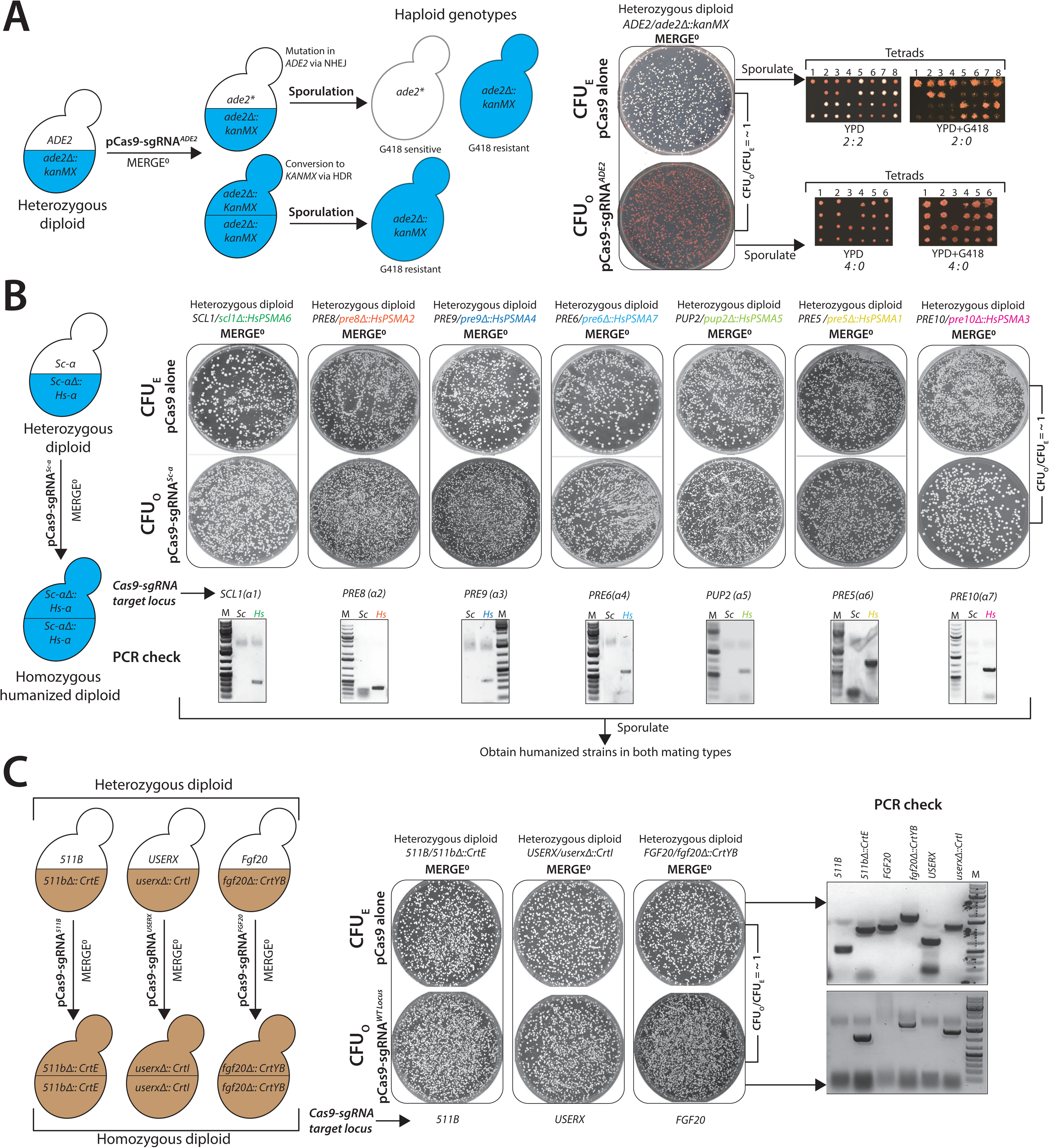
MERGE^0^ enables the efficient conversion of single heterozygous to homozygous loci. **(A)** A schematic of possible genotypes after the *ADE2* locus in a heterozygous diploid *ADE2*/*ade2Δ::kanMX* strain is subjected to a CRISPR-Cas9-mediated DSB followed by sporulation. The repair via NHEJ could mutate the locus (*ade2\**) or repair via HDR converts the locus to *ade2Δ::kanMX*, either scenario leads to a red colony phenotype. However, post sporulation, the prior scenario will show only 50% G418 resistant colonies compared to the latter outcome, which will exhibit 100% G418 resistance. Tetrad dissection of pooled cells from heterozygous diploid strains before **MERGE^0^** (**pCas9 alone**) or after **MERGE^0^** (**pCas9-sgRNA*^ADE2^***) shows that before **MERGE^0^** cells appear as 2 : 2 (red : white) and 2 : 2 (G418- resistant : G418-sensitive) phenotype. However, after **MERGE^0^,** cells show 4 : 0 (red : white) phenotype. Furthermore, all 4 spores in the post **MERGE^0^** strain show 4 : 0 G418 resistance. **(B)** Schematic shows CRISPR reagent targeted to one of the heterozygous alleles (wild type yeast gene) in a diploid strain permits efficient conversion to homozygous humanized loci. **MERGE^0^** efficiently converts wild-type yeast loci α proteasome genes to humanized loci (**CFU_O_/CFU_E_= ∼0**). PCR-based genotyping verified the conversion to the CRISPR-resistant humanized loci for every colony tested. **(C)** Schematic shows how CRISPR reagent targeted to the landing pad loci in a heterozygous diploid enables conversion to engineered carotenoid loci. **MERGE^0^** converts heterozygous landing pad loci (*511B, USERX,* and *FGF20*) to engineered carotenoid loci (**CFU_O_/CFU_E_ = ∼1)**. PCR-based genotyping shows heterozygous status of the loci before **MERGE^0^** and conversion to engineered carotenoid loci after **MERGE^0^**. Sporulation yields haploids of both mating types with only engineered loci.

**Figure S6.**
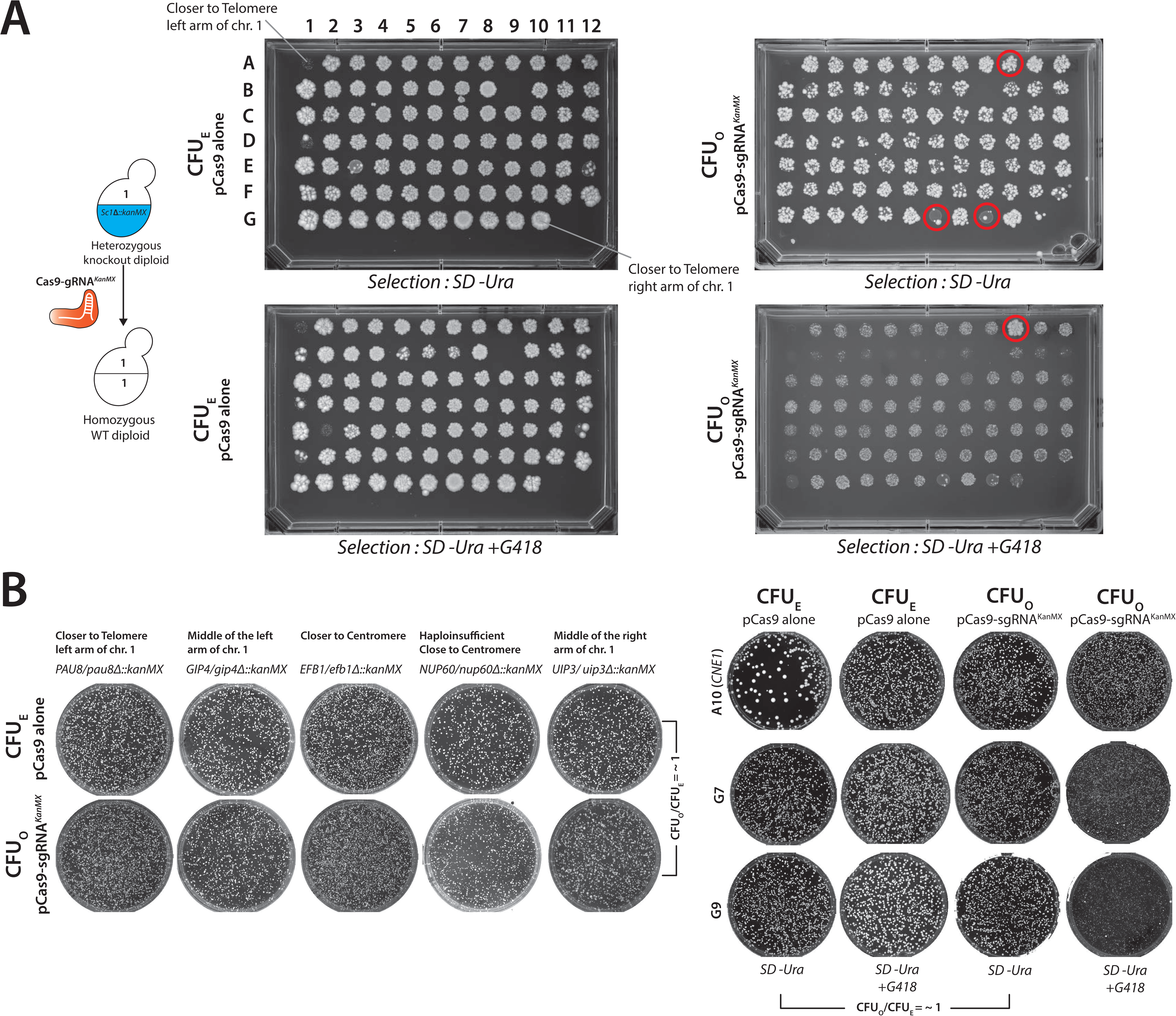
MERGE^0^ is efficient at converting many yeast heterozygous to homozygous loci independent of the position on Chromosome 1. **(A)** Schematic shows a CRISPR reagent (**pCas9-sgRNA*^KanMX^***) targeted to the KanMX allele in a hetKO strain converts the locus to homozygous wild-type. The hetKO strains were arrayed as genes close to the left-arm of a telomere through the centromere to the right arm of a telomere. The strains were transformed with **pCas9 alone** (**CFU_E_**) or with **pCas9-sgRNA*^KanMX^*** (**CFU_O_**) and spotted on either SD-Ura or SD-Ura with G418 selection. Every genotype shows spots with a similar density of colonies showing no lethality due to the CRISPR reagent (**CFU_O_/CFU_E_ = ∼1**). The spots on SD-Ura with G418 selection show G418 resistance in the case of **pCas9 alone** transformants. However, all but one (A10 - *CNE1*) strain transformed with **pCas9-sgRNA*^KanMX^*** lost the *KanMX* cassette resulting in G418 sensitivity. **(B) MERGE^0^** mediated conversion of loci was tested on several unique hetKO genotypes on Petri plates, demonstrating agreement with 96-well spot assays. All tested strains show no lethality due to the CRISPR reagent (**CFU_O_/CFU_E_ = ∼1**). Ambiguous spots on a 96-well plate (A10, G7 and G9) were further analyzed on Petri plates. G7 and G9 spots show no lethality due to CRISPR reagent (**CFU_O_/CFU_E_ = ∼1**) and loss of *KanMX* cassette. While the A10 (*CNE1*) strain is resistant to the CRISPR reagent, the cells retain G418 resistance. It is unclear why this behavior occurs.

**Figure S7.**
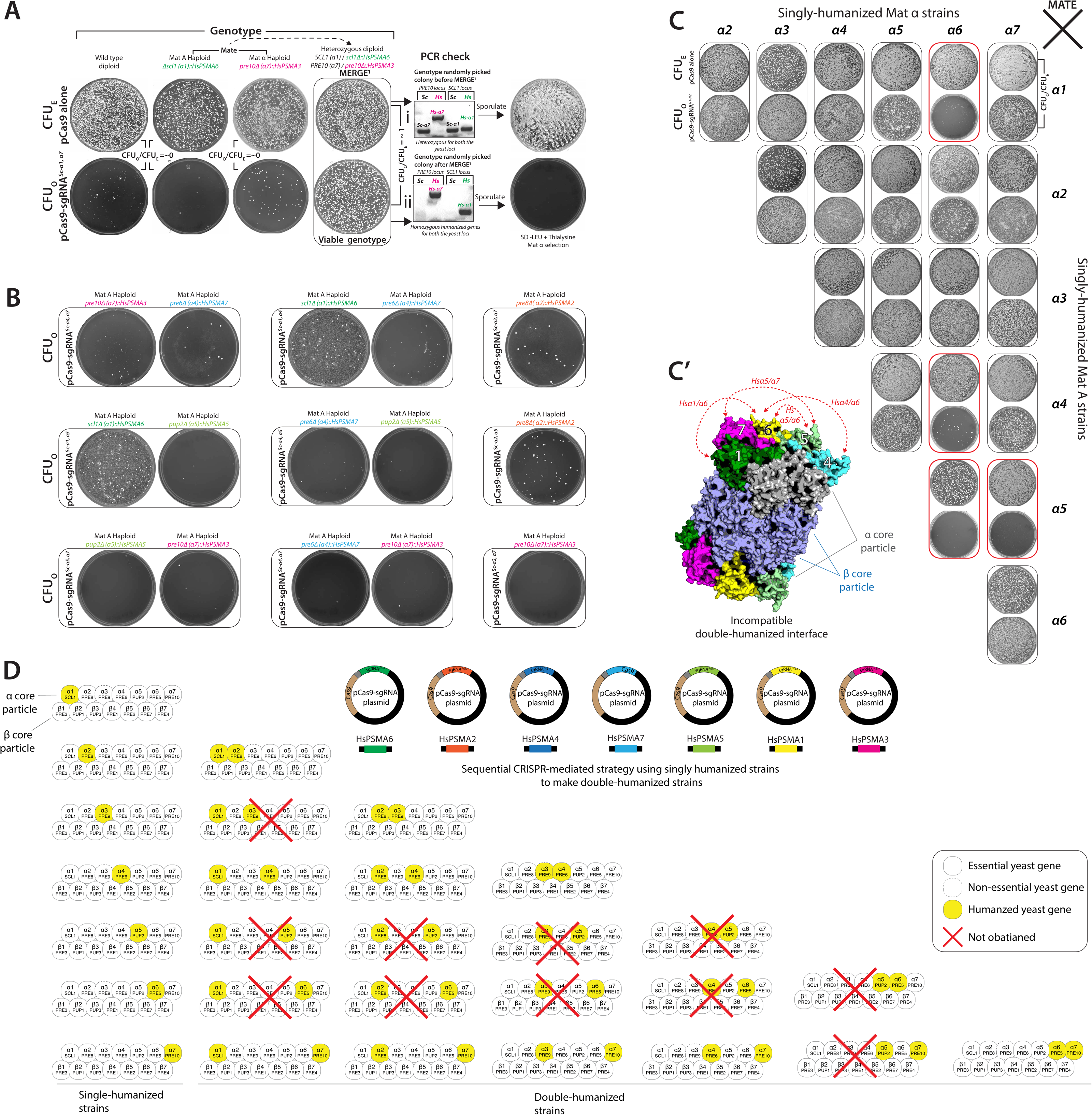
MERGE^0^ is efficient at simultaneously combining two heterozygous to homozygous loci. **(A)** Double-sgRNA CRISPR reagent (**pCas9-sgRNA^Sc^*^α1,α7^***) targeted to two wild type yeast loci (proteasome *α*1 and *α*7 genes) quantifies the efficiency of **MERGE^1^**. The transformation of **pCas9-sgRNA^Sc^*^α1,α7^*** in the wild type, and single-humanized Hs*α*1 or Hs*α*7 strains is lethal, suggesting ON-target activity (**CFU_O_/CFU_E_ = ∼0**). However, the transformation of **pCas9-sgRNA^Sc^*^α1,α7^*** in diploid heterozygous humanized strain shows no lethality (**CFU_O_/CFU_E_ = ∼1**). PCR-based genotyping of surviving colonies after **MERGE^1^** shows conversion of both yeast to the humanized loci compared to before **MERGE^1^. (B)** All tested single-humanized strains transformed with double-sgRNA CRISPR reagents show lethality, demonstrating that selection only allows survival of viable paired genotypes. **(C)** Mating each single-humanized strain of opposite mating-types (obtained via **MERGE^0^**) provided 21 heterozygous genotypes to test **MERGE^1^**. Heterozygous diploid humanized strains transformed with double-sgRNA CRISPR reagents specific to each yeast allele allow testing viability of all paired-humanized genotypes (**CFU_O_**). If the genotype is permitted, the strategy enables survival similar to the pCas9 alone transformation (**CFU_E_**). While the majority of combined double-humanized genotypes are viable (**CFU_O_**/**CFU_E_ = ∼1**), specific genotypes (*Hsα1α6*, *Hsα4α6*, *Hsα5α6* and *Hsα5α7*) were inviable, suggesting incompatible genotypes. Other combinations, such as those involving the genotypes with Hs*α*2, showed a slow-growth phenotype. **(C’)** Yeast proteasome structure (PDB-1RYP) with *α* and β cores show the humanized *α* subunits that are incompatible as paired genotypes. Except for *α5* and *α6*, the subunits are missing neighboring interactions within the α core. **(D)** A sequential strategy using single-humanized strains acquired several pairwise combinations of human α subunits. However, several other genotypes could not be obtained, revealing the drawback of the method. **MERGE^1^** is far more efficient in identifying incompatible paired genotypes.

**Figure S8.**
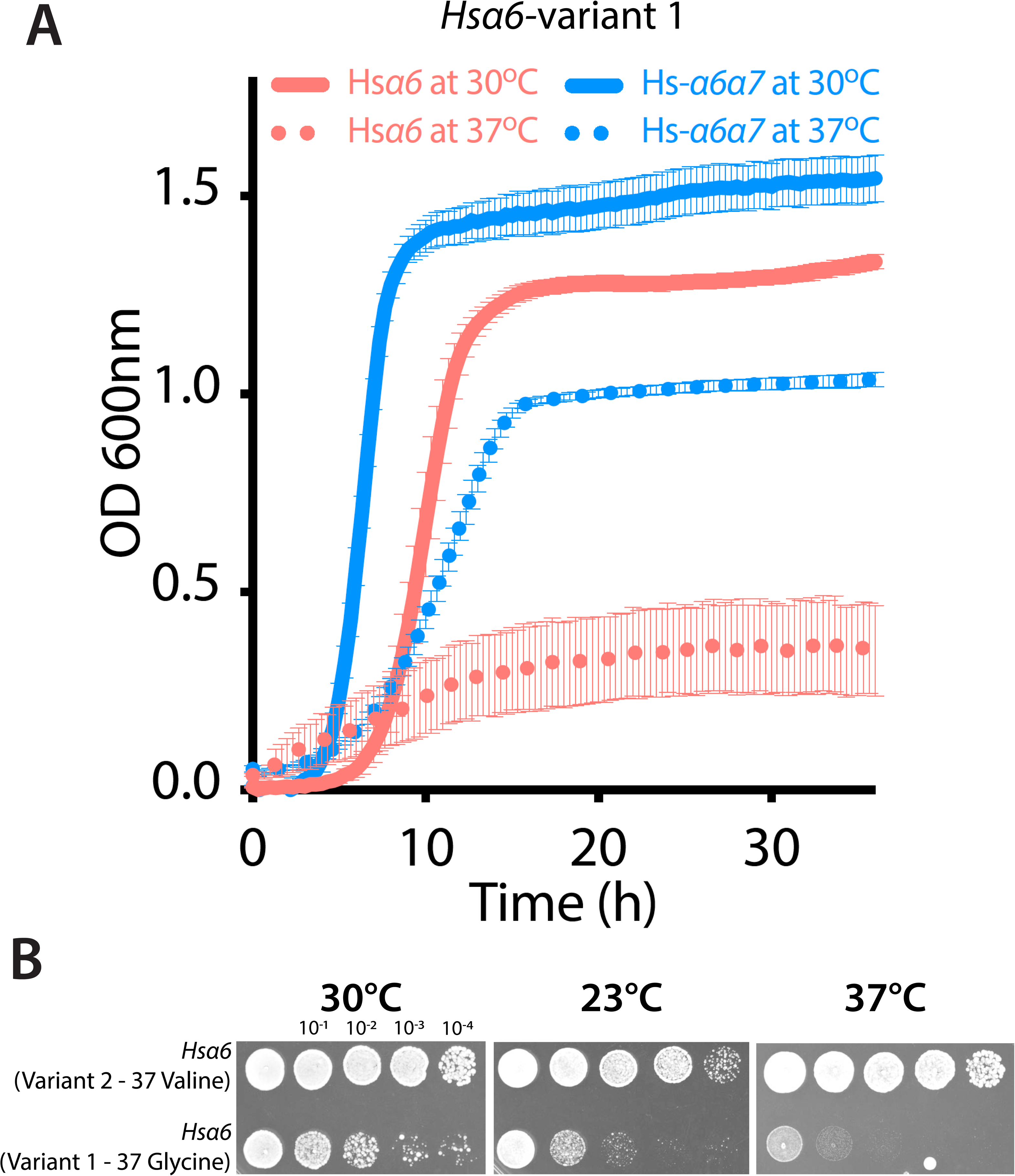
The variant-specific phenotype of single-humanized Hs*α6* (*PSMA1*) yeast strain. **(A)** A growth assay performed on a single-humanized *Hsα6* strain (Variant 1, amino acid residue 37>Glycine) shows a temperature-sensitive (TS) phenotype at 37°C (dotted red line) compared to growth at 30°C (solid red line). The combined genotype of Hs*α*6,*α*7 partially rescues the TS phenotype at 37°C (dotted blue line) compared to the growth at 30°C (solid blue line). **(B)** Spotted dilutions of single-humanized *Hsα6* strains (Variant 1, amino acid residue 37>Glycine) and (Variant 2, amino acid residue 37>Valine) show Variant 2 with no growth defect at 30°C, 23°C and 37°C compared to Variant 1.

**Figure S9.**
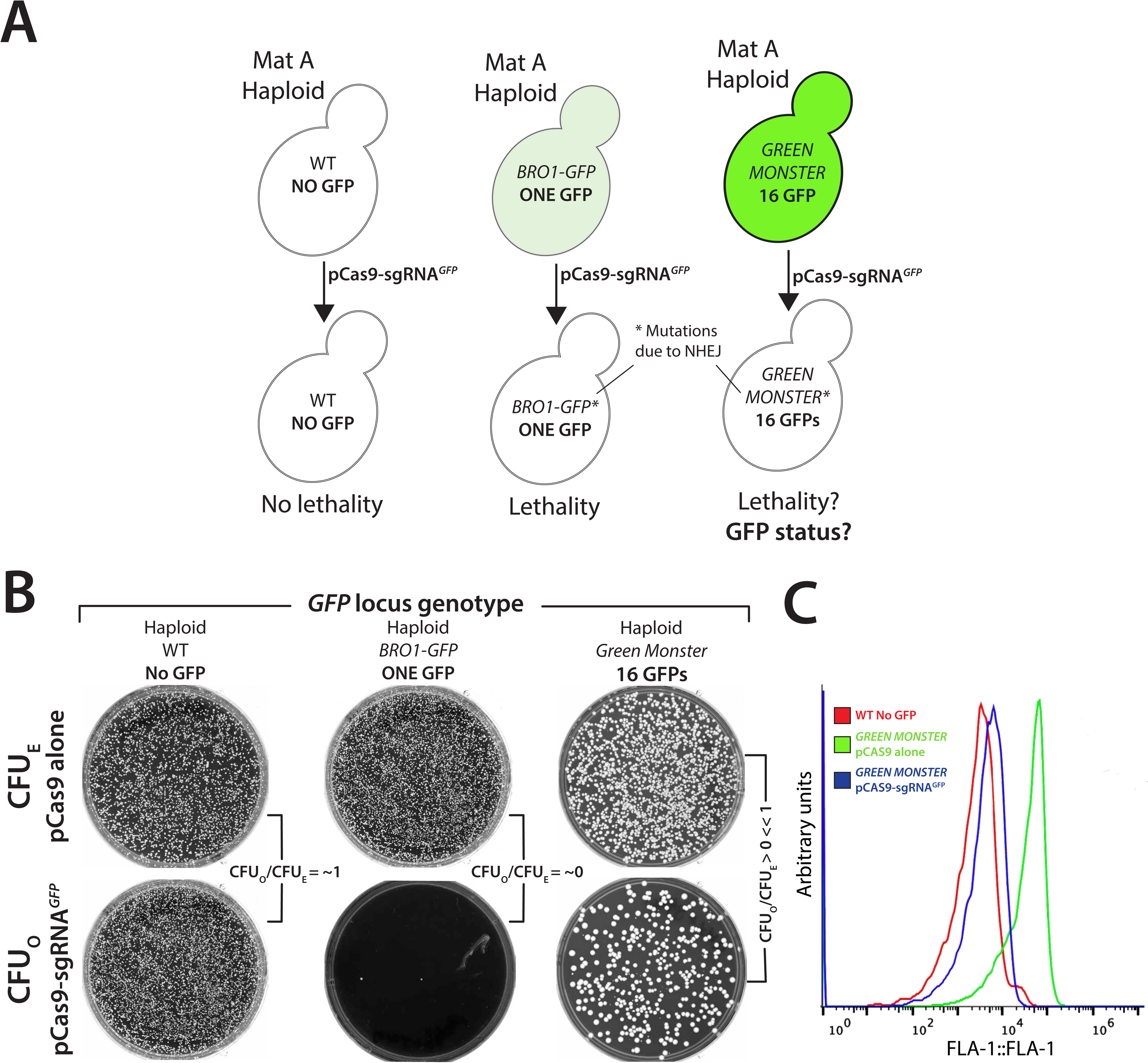
MERGE is scalable to target multiple yeast loci simultaneously. **(A)** The schematic shows a CRISPR-based strategy targeting one or many genetic loci in yeast. Transformation of a CRISPR reagent (**pCas9-sgRNA^GFP^**) targeting a GFP cassette in a Green Monster strain tests the scalability. Transformation in the wild type (no GFP) strain should show no lethality, whereas strains harboring single or multiple GFPs should show a lethal phenotype. **(B)** The **pCas9-sgRNA^GFP^** transformation in wild type yeast showed viable cells (**CFU_O_/CFU_E_ =∼1**). In contrast, transformation in a strain harboring a single GFP cassette was lethal. While the Green Monster strain (GFP monster, 16 GFP cassettes) showed more viable cells than the single-GFP strain upon transformation with **pCas9-sgRNA^GFP^**. **(C)** Using flow cytometry, the pooled mixture of yeast cells that survive **pCas9-sgRNA^GFP^** showed significantly reduced GFP expression compared to the pCas9 alone transformed cells, suggesting that nearly every GFP cassette was successfully targeted and mutated via NHEJ.

**Figure S10.**
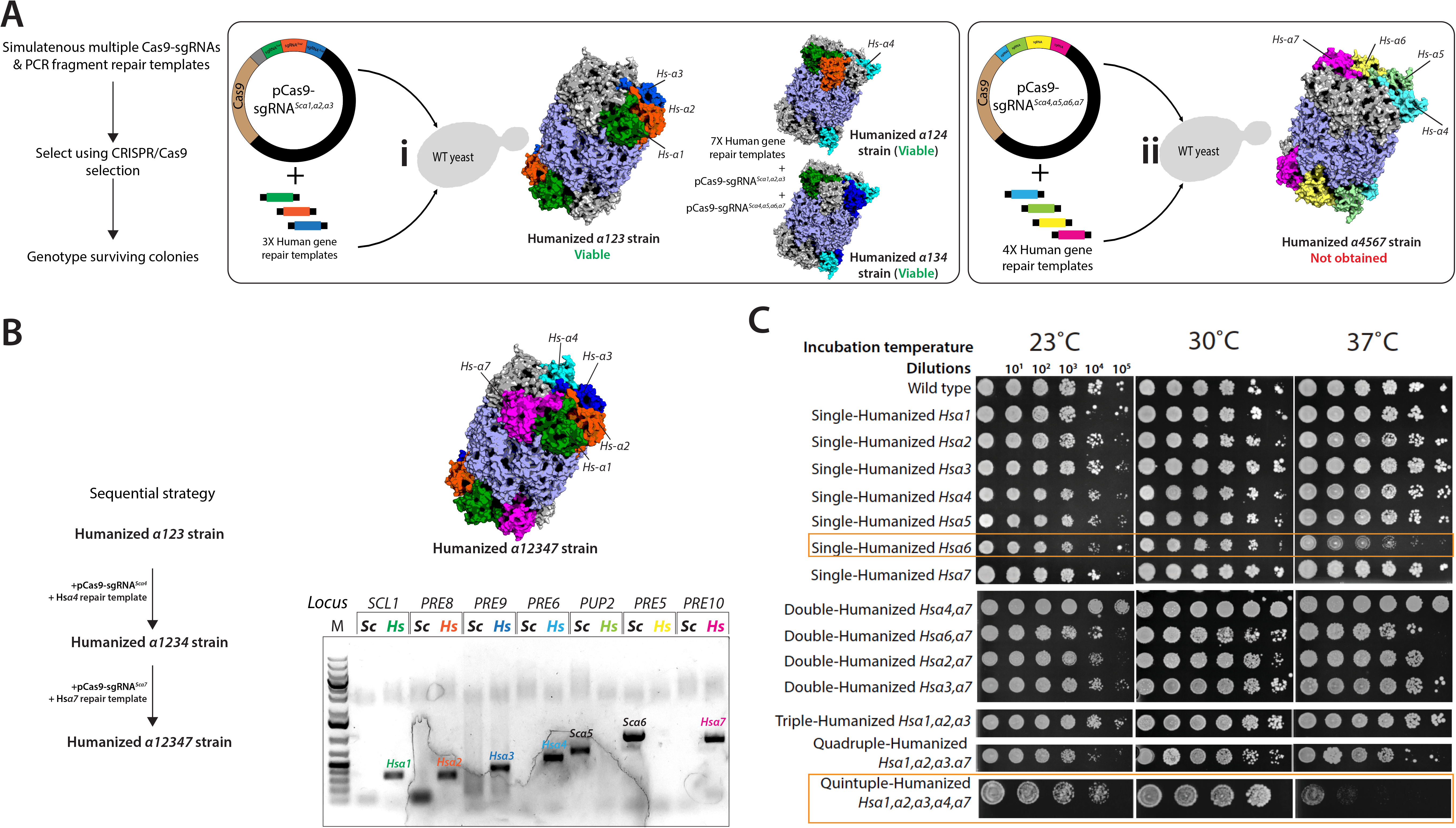
Scheme for obtaining triple-, quadruple-, quintuple- and sextuple-humanized α-subunit strains *via* a sequential strategy. **(A)** Two multiple-sgRNA CRISPR plasmids were constructed, one expressing sgRNAs targeting yeast α subunits 1, 2, and 3 (**pCas9-sgRNA*^Scα1,α2,α3^***), and the other expressing sgRNAs targeting α subunits 4, 5, 6, and 7 (**pCas9-sgRNA*^Scα4,α5,α6,α7^***). They were transformed individually with the respective human gene repair DNA. A single colony with triple-humanized *Hs-α1α2,α3* (**i**) was obtained. However, no clone for a quadruple humanized *Hs-α4α5,α6,α7* (**ii**) could be obtained. A similar strategy also generated triple-humanized *Hs-α1α2,α4* and *Hs-α1α3,α4* genotypes. **(B)** Sequential strategy generated a quintuple-humanized *Hs-α1α2,α3*,*α4,α7* strain confirmed via locus-specific PCR showing the presence of human and the absence of yeast loci. Sanger sequencing and whole-genome sequencing confirmed the genotype. **(C)** Spotted dilutions of double-, triple-, quadruple-, quintuple- and sextuple humanized *α* proteasome strains at 23°C, 30°C and 37°C show singe-humanized Hs-*α6* (variant 1) and quintuple-humanized *Hs- α1,α2,α3*,*α4,α7* strain with a temperature-sensitive growth phenotype.

**Figure S11.**
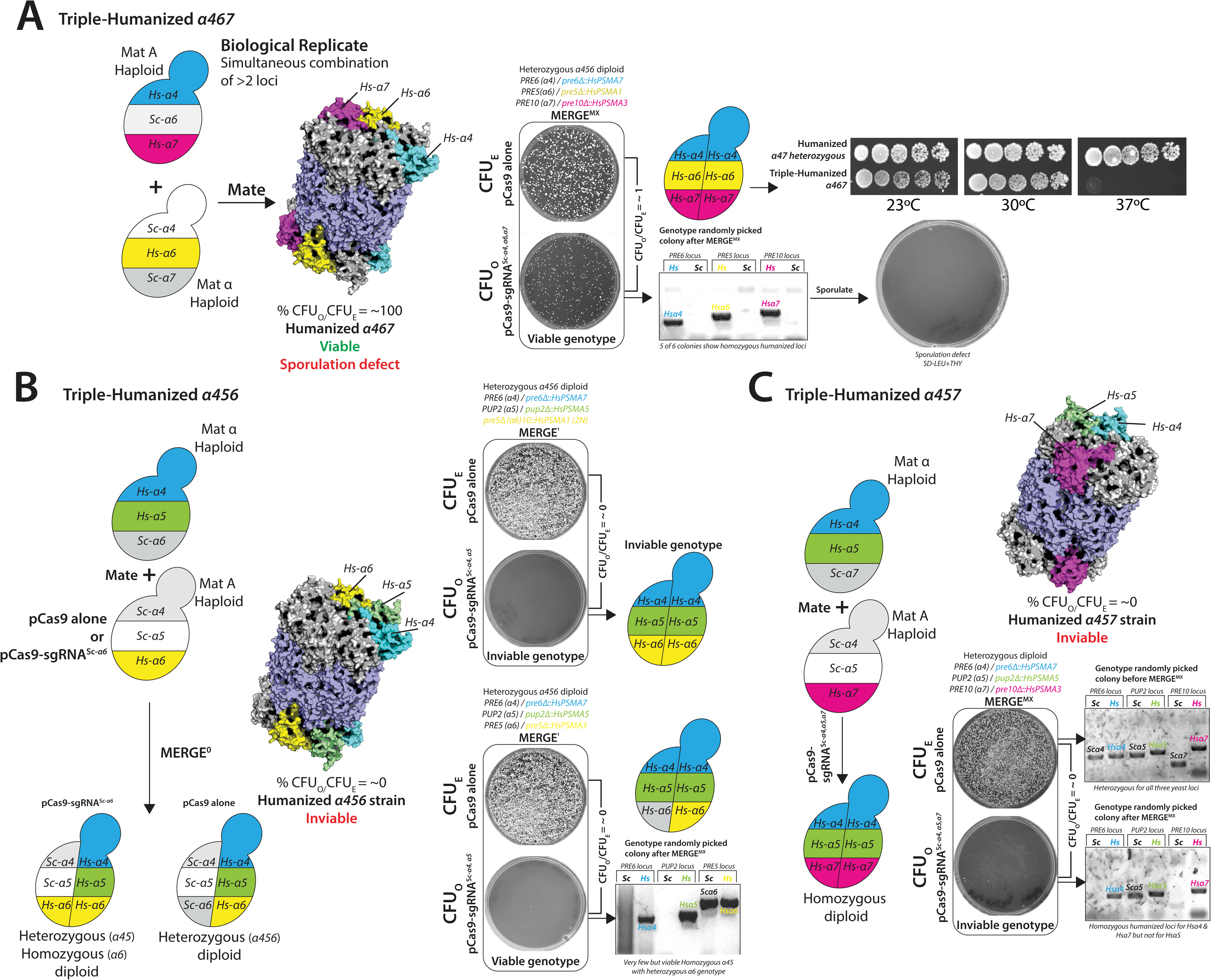
MERGE tests the combination of several triple-humanized yeast α-subunit genotypes. (A) **MERGE^MX^** (**Mate and multiplex**) strategy combined 3 loci after mating a strain with 2 humanized loci (Hs*α*4,*α*7, Mat A) with a strain carrying 1 humanized locus (Hs*α*6, Mat α) (biological replicate of data shown in Figure 4E). The transformation of **pCas9-sgRNA*^Scα4,α6,α7^*** in a mated mix selected a diploid triple-humanized strain (**CFU_O_/CFU_E_ = ∼1**). PCR-based genotyping of several colonies after **MERGE^MX^** shows conversion of all yeast to the humanized loci (5 of 6). Spotted dilutions of humanized Hs-*α*4,*α*7 and Hs-*α*4,*α*6,*α*7 strains at 23°C, 30°C and 37°C show the triple-humanized strain with a temperature-sensitive phenotype at 37°C. The strain does not yield viable haploid spores observed as no growth on haploid-specific selection media (SD-LEU+Thialysine). (B) Double-humanized *Hs-α4α5* (haploid Mat α) and single-humanized *Hs-α6* (haploid Mat A, harboring a **pCas9 alone** or **pCas9-sgRNA*^Sc-α6^*** vector) strains were mated, generating two distinct heterozygous diploid strains, one heterozygous for three loci and another for two with homozygous *Hs-α6* locus. Each strain was transformed with double-sgRNA CRISPR plasmid (**pCas9-sgRNA*^Scα4,α5^***) to perform **MERGE^1^**. The transformation caused lethality in both strains (**CFU_O_/CFU_E_ =∼0**). However, a few survivors in the context of heterozygous *Hs-α6* locus showed the conversion of two loci (*Hs-α4* and *Hs-α7*), suggesting an incompatible triple-humanized genotype but viable double-humanized genotype harboring a heterozygous yeast-human *α6* locus. **(C)** Double-humanized haploid *Hs-α4α5* (Mat α) and *Hs-α7* (Mat A) strains were mated. The resulting heterozygous diploid strain was transformed with a triple-sgRNA CRISPR reagent (**pCas9-sgRNA*^Scα4,α5,α7^***). The transformation caused lethality with few survivors (**CFU_O_/CFU_E_ =∼0**). Genotyping of the survivors showed the conversion of two 2 of 3 loci (*Hs-α4* and *Hs-α7*), suggesting an incompatible triple-humanized genotype.

**Figure S12.**
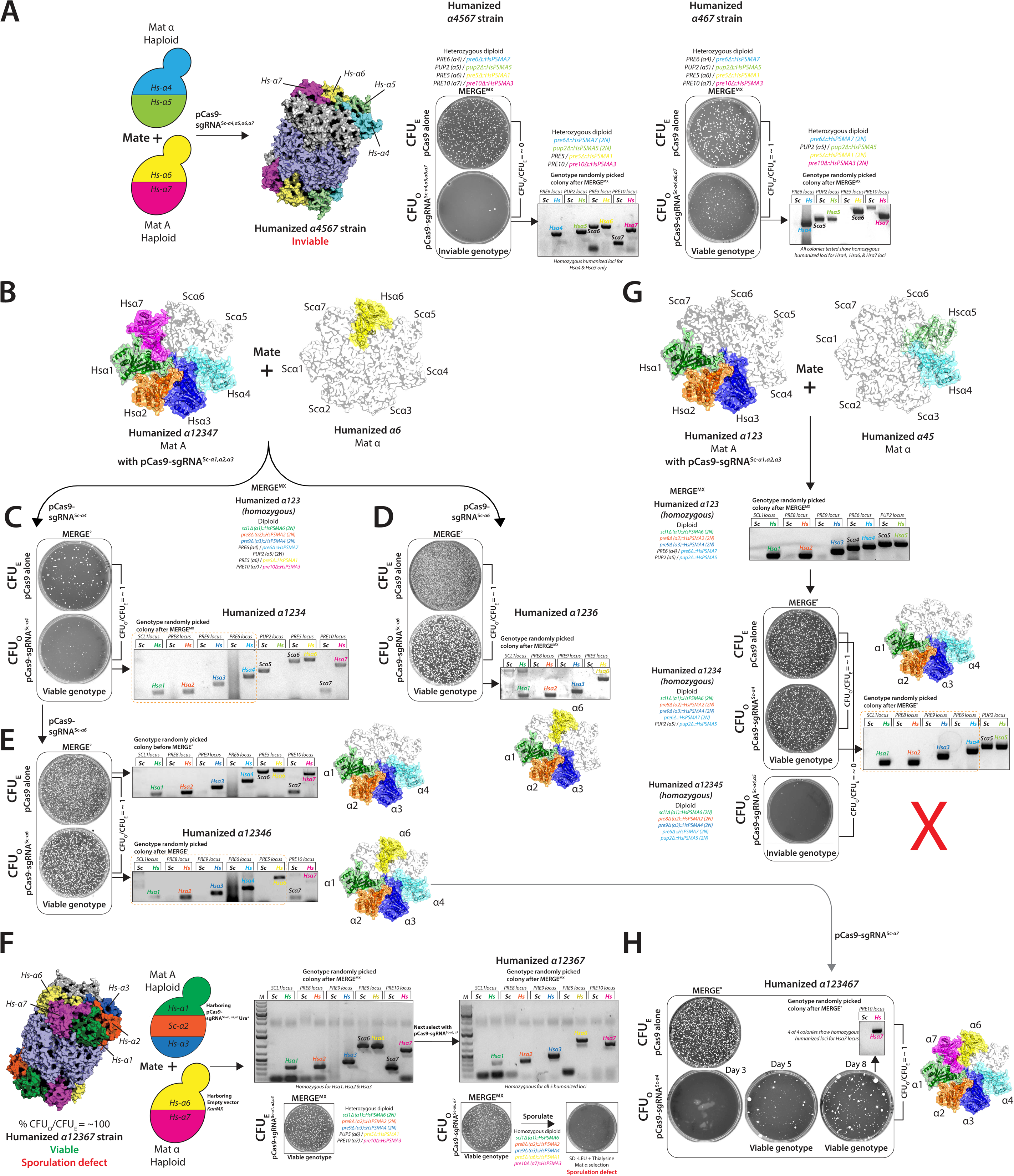
MERGE efficiently tests several higher-order combinations (>3) of humanized yeast α-proteasome core. (**A**) Double-humanized haploid *Hs-α4α5* (Mat α) and *Hs-α6α7* (Mat α) strains were mated. The resulting heterozygous diploid strain was transformed with quadruple-sgRNA CRISPR reagent (**pCas9-sgRNA*^Scα4,α5,α6,α7^***) to perform **MERGE^MX^**. The transformation caused lethality with few survivors (**CFU_O_/CFU_E_ =∼0**). The survivors showed the conversion of only 2 of 4 loci (*Hs-α4* and *Hs-α5*), suggesting an incompatible quadruple-humanized genotype. However, the transformation of triple-sgRNA CRISPR reagent (**pCas9-sgRNA*^Scα4,α6,α7^***) in this strain was viable (**CFU_O_/CFU_E_ =∼1**). The genotyping of randomly picked colonies showed the conversion of 3 yeast to human loci (*Hs-α4, Hs-α6,* and *Hs-α7*), suggesting a triple-humanized genotype is viable with a heterozygous yeast-human *α5* locus. **(B)** Mating quintuple-humanized haploid *Hs-α1α2α3α4α7* [Mat A, also harboring a triple-sgRNA CRISPR reagent (**pCas9-sgRNA*^Scα1,α2,α3^***)] and single-humanized *Hs-α6* (Mat α) strains simultaneously combined three yeast to humanized *Hsα1, Hsα2, Hsα3* loci in a diploid yeast (**CFU_O_/CFU_E_ =∼1**). The strain harbors heterozygous yeast-human alleles for *α4, α6,* and *α7* loci. Transformation of the diploid strain with **(C) pCas9-sgRNA*^Scα4^*** or **(D) pCas9-sgRNA*^Scα6^*** generated viable quadruple-humanized strains (Hs-*α1α2α3α4* and Hs- *α1α2α3α6*) (**CFU_O_/CFU_E_ =∼1**). Genotyping of randomly picked colonies confirmed the homozygosity of the loci. **(E)** Using the quadruple-humanized strain *Hs-α1α2α3α4* as a background, the transformation of **pCas9-sgRNA*^Scα6^*** yielded a viable quintuple-humanized (Hs- *α1α2α3α4α6*) genotype (**CFU_O_/CFU_E_ =∼1**) confirmed using locus-specific PCR. **(F)** Double-humanized haploid *Hs-α6α7* (Mat α) and triple-humanized *Hs-α1,α2,α3* [Mat A, also harboring a triple-sgRNA CRISPR reagent (**pCas9-sgRNA*^Scα1,α2,α3^***)] strains were mated. **MERGE^MX^** simultaneously combined three yeast to humanized *Hsα1, Hsα2, and Hsα3* loci in the diploid background (**CFU_O_/CFU_E_ =∼1**). The next transformation, using **pCas9-sgRNA*^Scα6,α7^*** also homozygosed the *Hsα6* and *Hsα7* loci, generating a quintuple-humanized strain (**CFU_O_/CFU_E_ =∼1**). However, post sporulation, the mix plated on Mat α selection failed to produce any viable humanized haploid strains. **(G)** Triple-humanized *Hs-α1α2α3* [haploid Mat A, also harboring a triple-sgRNA CRISPR reagent (**pCas9-sgRNA*^Scα1,α2,α3^***)] and double-humanized *Hs-α4α5* (haploid Mat α) strains were mated generating a heterozygous diploid strain for 5 loci. The resulting diploid strain simultaneously combined three yeast to humanized *Hsα1, Hsα2, and Hsα3* loci while harboring two yeast-human heterozygous loci (*α4* and *α5)* (**CFU_O_/CFU_E_ =∼1**). The subsequent **MERGE^0^** strategy (using **pCas9-sgRNA*^Scα4^***) generated a viable quadruple-humanized strain (*Hs-α1α2α3α4*) with heterozygous yeast-human *α5* locus (**CFU_O_/CFU_E_ =∼1**). The locus-specific PCR confirmed the homozygosity for most of the colonies. However, the subsequent CRISPR selection (using **pCas9-sgRNA*^Scα5^***) did not produce a viable strain (**CFU_O_/CFU_E_ =∼0**), suggesting an incompatible quintuple-humanized (*Hs-α1α2α3α4α5*) genotype. **(H)** Using the quintuple-humanized diploid strain *Hs-α1α2α3α4α6* harboring a yeast-human heterozygous *α7* locus as a background, the transformation of **pCas9-sgRNA*^Scα7^*** yielded a viable sextuple-humanized (Hs-*α1α2α3α4α6α7*) genotype (**MERGE^0^, CFU_O_/CFU_E_ =∼1**) after incubation for 8 days. The genotype of the viable colonies confirmed the homozygosity and humanization of the *α7* locus.

**Figure S13.**
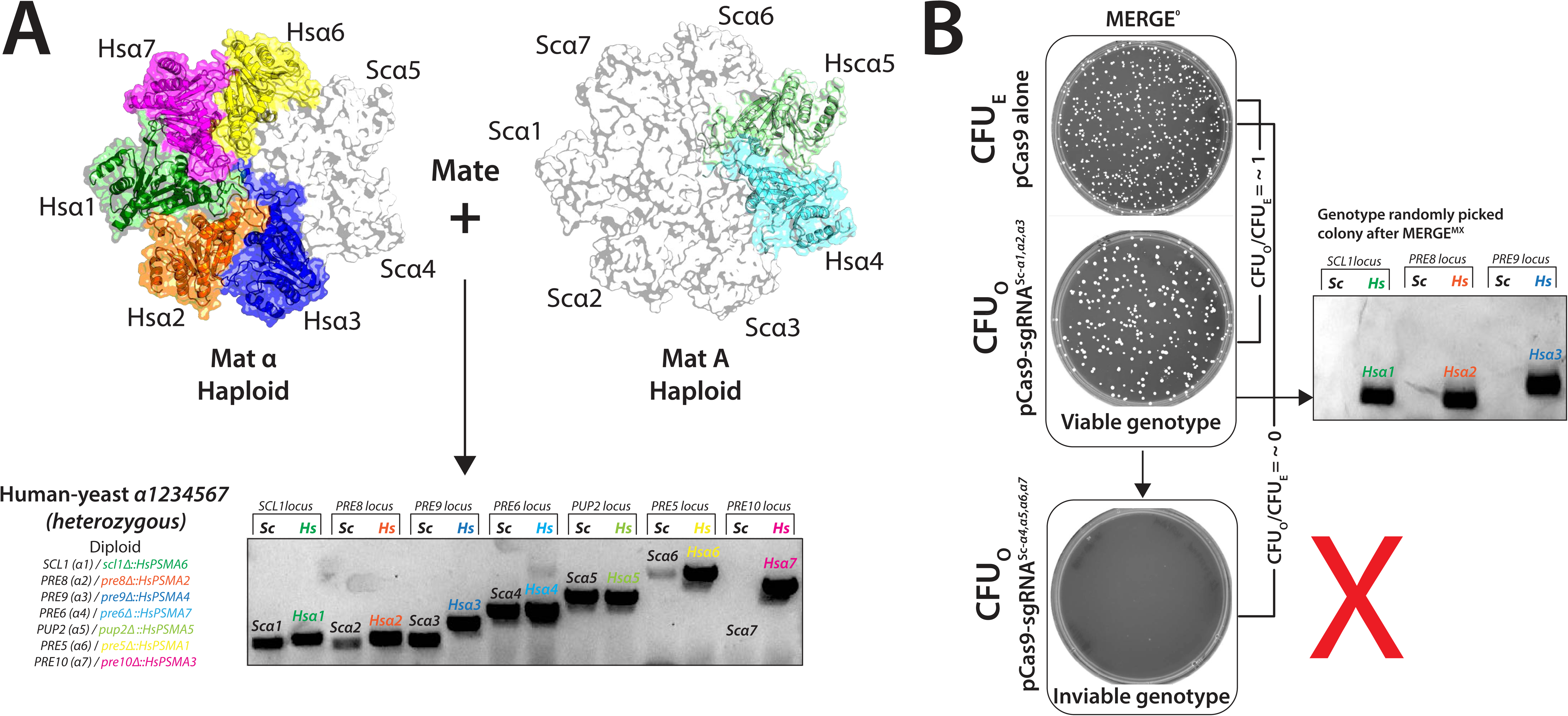
MERGE enables the verification of the entire human α-proteasome core in yeast. **(A)** Mating quintuple-humanized haploid *Hs-α1α2α3α6α7* (Mat α) and double-humanized *Hs-α4α5* (Mat A) strains generated a diploid strain harboring yeast-human heterozygous alleles for 7 *α-*proteasome loci. Locus-specific PCR confirmed the heterozygous nature of the loci. **(B)** The resulting diploid strain was transformed with a triple-sgRNA CRISPR reagent (**pCas9-sgRNA*^Scα1,α2,,α3^***), resulting in a viable genotype homozygous for triple-humanized Hs-*α1α2α3* loci (**CFU_O_/CFU_E_ =∼1**) with yeast-human heterozygous alleles for *α4, α5, α6,* and *α7* loci. The subsequent transformation of a quadruple-sgRNA CRISPR reagent (**pCas9-sgRNA*^Scα4,α5,α6,α7^***) in the diploid strain was lethal, suggesting an incompatible genotype (**CFU_O_/CFU_E_ =∼0**).

**Figure S14.**
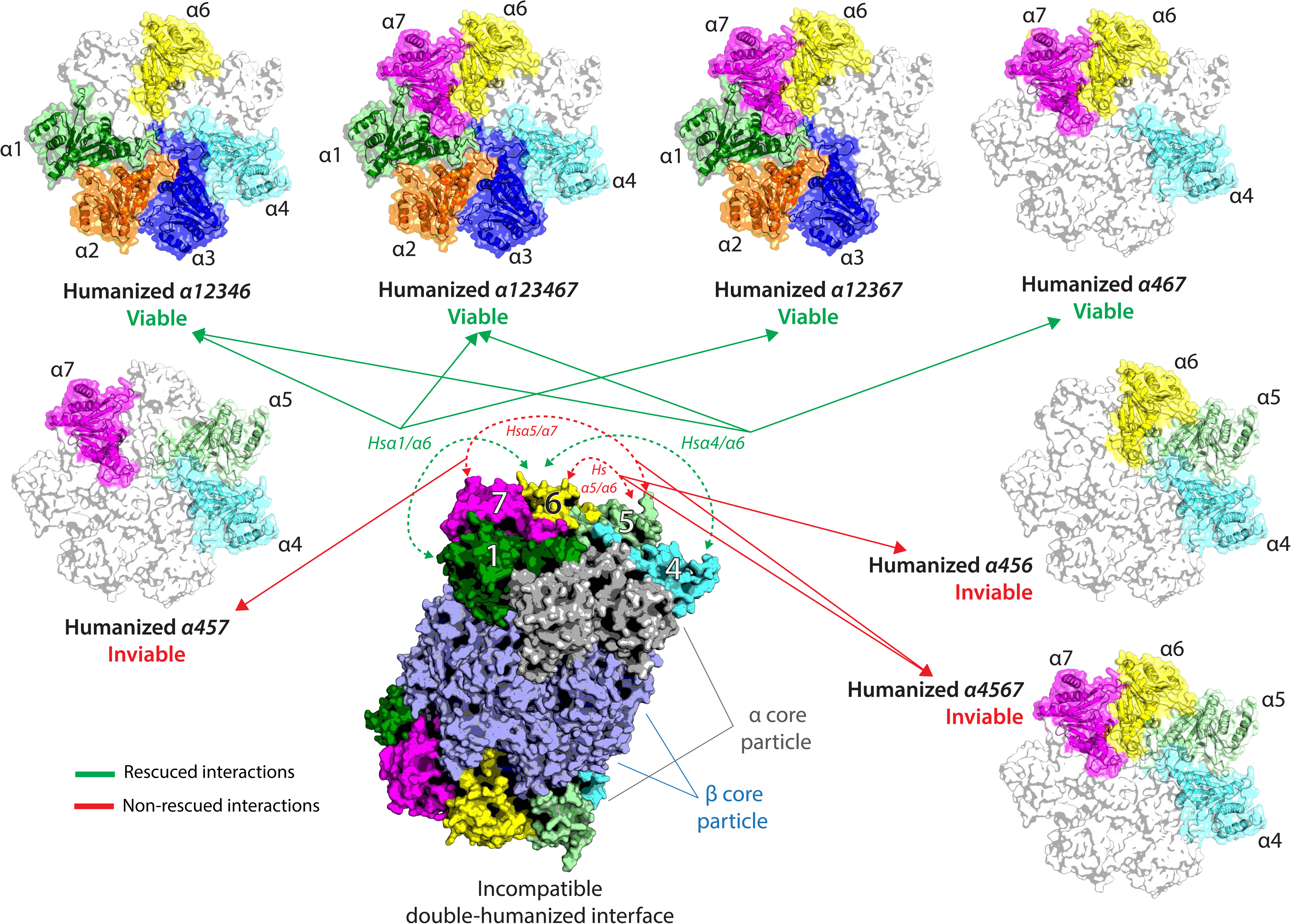
Inviable double-humanized genotypes are viable when neighboring interactions are restored. Inviable double-humanized genotypes, *Hsα1/α6* and *Hsα4/α6,* become viable when the neighboring interactions are restored in higher-order humanized strains (indicated as green lines). However, incompatible paired genotypes comprising *Hsα5/α6* and *Hsα5/α7* could not be rescued (marked as red lines). Proteasome core structures were generated using Pymol and PDB-1RYP. Colored structures show humanized *α* yeast subunits.

**Figure S15.**
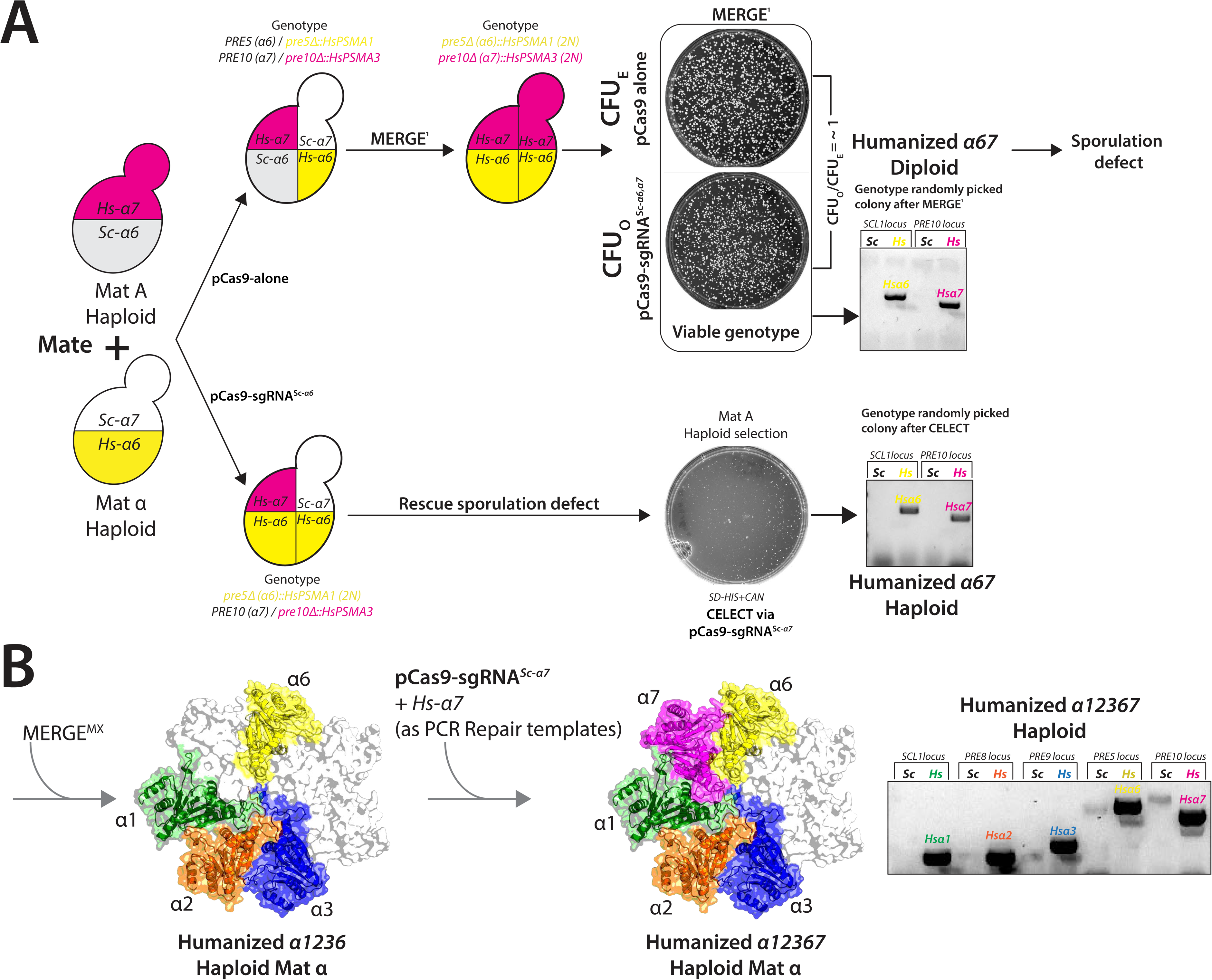
Strategies to generate viable humanized haploid yeast strains when the corresponding diploids manifest sporulation defects. **(A)** Mating single-humanized *Hs-α6* (Mat α, with **pCas9 alone** or **pCas-sgRNA*^Sc-α6^***) and *Hs-α7* (Mat A) strains generated two diploid strains; one harboring yeast-human heterozygous allele at *α*6 locus and another with homozygous humanized *α6* locus. **MERGE^0^**(using strategy **pCas-sgRNA*^Sc-α7^***) converted the heterozygous yeast-human *α7* locus to a homozygous human allele at ∼100% efficiency (**CFU_O_/CFU_E_ =∼1**). Locus-specific PCR confirmed the humanized homozygous alleles. However, the diploid strains cannot sporulate. However, keeping the *α7* locus as heterozygous rescues the sporulation defect. The haploid-specific selection (SD-HIS+CAN; Mat A) allowed haploid yeast to grow, followed by **CELECT** (using **pCas-sgRNA*^Sc-α7^***) or simply genotyping the haploids to identify homozygous double-humanized Hs-*α6α7* genotype. **(B)** The diploid quintuple-humanized Hs-*α1α2α3α6α7* strain shows a sporulation defect as in **Figure S12F**. However, using a sequential CRISPR strategy, a haploid quadruple-humanized Hs-*α1α2α3α6* strain enables the generation of haploid quintuple-humanized Hs-*α1α2α3α6α7* strain to be used for subsequent **MERGE** experiments.

**Figure S16.**
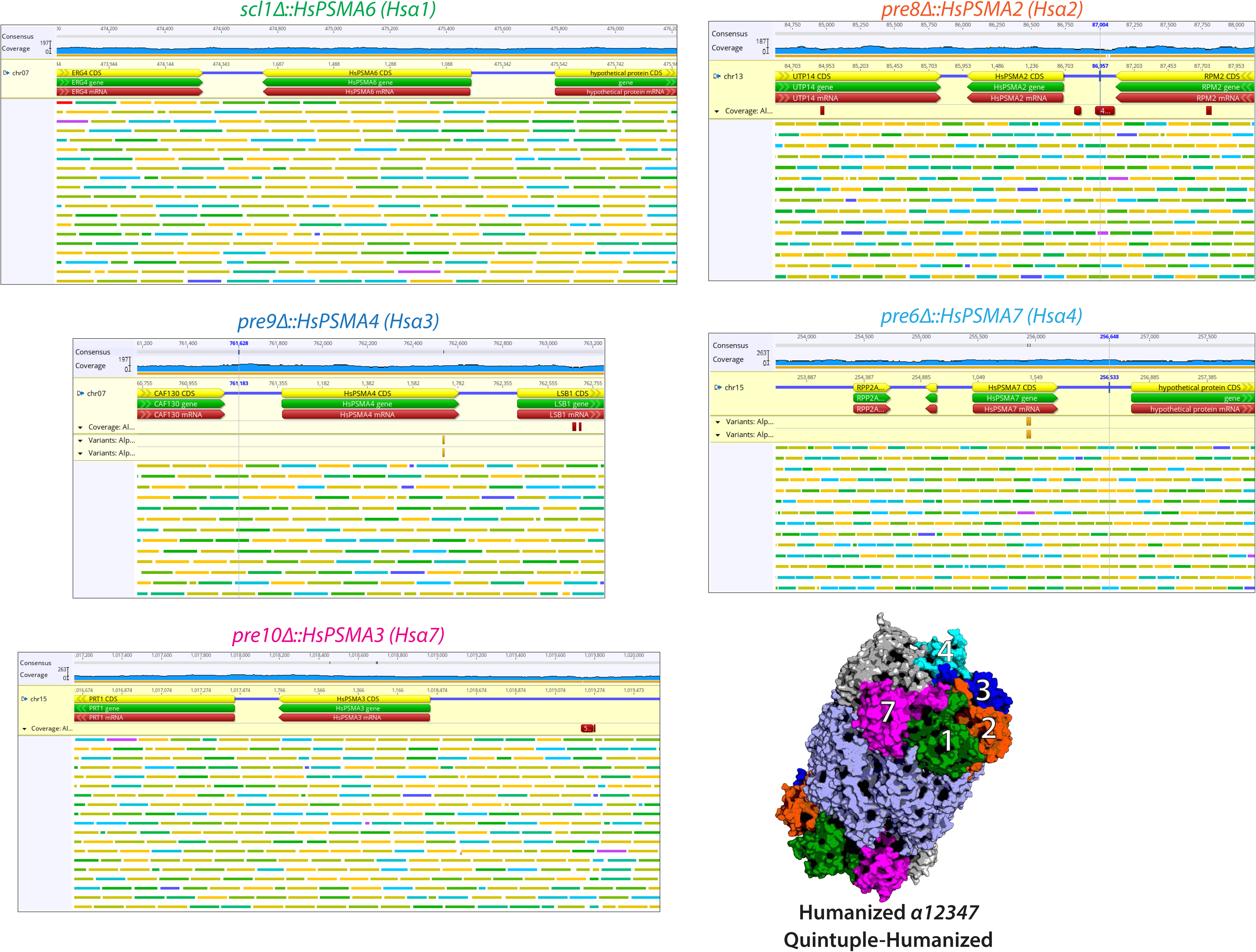
Whole-genome sequencing (WGS) of quintupled-humanized *Hs-α1α2α3α4α7* strain shows humanized loci alignment within the yeast genome. WGS analysis of quintupled-humanized *Hs-α1α2α3α4α7* shows humanized loci aligned to a reference S288C genome with replaced human coding sequences. The human genes reside on different yeast chromosomes, replacing their corresponding native yeast loci (Chr. 7, 13, and 15). Green lines indicate the read lengths with a 200 base range, yellow is below, and blue is above the range.

## Materials & methods

### 1. Targeting sgRNA design

CRISPR-Cas9 targeting sequences consist of a 20 bp recognition sequence preceding an ‘NGG’ sequence motif (PAM). To design our targeting sequences, we used the built-in guide RNA design tool in version 11 of the Geneious software^36^, using a recent version of the yeast genome (available from http://www.yeastgenome.org/strain/S288C/overview) to screen for possible off-target sequences^13, 14^. We chose two high-scoring guide sequences to target each gene, requiring that they be near the 5’ end of the gene so any NHEJ repair would likely result in an early frameshift. The guides were ordered as complementary oligos with overhangs according to the sgRNA template described in the Yeast Tool Kit^15^ (**Table S1**).

### 2. Plasmid construction using Yeast Tool Kit (YTK)

Entry vectors for each guide RNA sequence were constructed by the BsmbI-mediated Golden Gate (GG) assembly (Thermo) into plasmid pYTK050 from the Yeast ToolKit^15^. The targeting sequences were ordered (IDT) as two single-stranded DNA oligos with a complementary region and unique overhangs according to the ‘sgRNA’ template described in Lee, ME *et al*^15^. Briefly, the two oligos for each guide were annealed by slow cooling from 95°C to 4°C (1-5 hrs) in a thermal cycler, and 1μl of the annealed product was added to the entry vector GG reaction. Entry Vectors were sequence-verified using custom primers.‘Transcriptional unit’ (TU) vectors (pYTK095) were constructed by a BsaI-mediated GG assembly (Thermo). The entry vector for a particular guide RNA was combined with a left connector part plasmid (ConLX) and right connector part plasmid (ConRX) into the AmpR-ColE1 (pYTK095) backbone plasmid^13, 14^. To create single-sgRNA TU vectors, we assembled the sgRNA entry vector with left connector 1 (ConL1, pYTK003) and right connector E (ConRE, pYTK072). To create TU vectors for the multi-sgRNA knockout, an entry vector was combined with the appropriate connectors to enable proper subsequence knockout vector assembly (i.e. a triple-sgRNA vector would have one TU vector with ConL1 and R2, one with ConL2 and R2, and the third with ConL3 and RE). ‘Knockout’ (KO) vectors were constructed by BsmbI-mediated GG assembly (Thermo). The appropriate TU vectors were assembled into the CEN6-URA3 backbone and the Cas9-TU1 vector to create a KO vector. Each GG reaction was performed in a 10μl volume, with approximately 20 fmol of each starting DNA molecule, along with 0.5μl each of the appropriate restriction enzyme, 0.5μl of T7 DNA ligase (NEB), and 1μl of 10X T4 ligase buffer (NEB). A thermocycler was used to cycle between 16°C and 37°C each for (5 minutes). 3μl of the reaction volume was transformed into competent DH5α *E. coli* and plated on appropriate selective media for the backbone (i.e. chloramphenicol for entry vectors, ampicillin for TU vectors, and kanamycin for KO vectors). Each backbone plasmid is a ‘GFP-dropout’ vector, so correct clones were selected by screening for non-fluorescent colonies when viewed by blue light and verified by sequencing.

### 3. Cas9-TU1

The Cas9-TU1 vector was constructed by BsaI GG assembly of YTK parts pYTK002 (ConLS), 011 (PGK1 promoter), 036 (Cas9 coding sequence), 055 (ENO2 terminator, 067 (ConR1), and 095 (AmpR-ColE1 GFP dropout backbone).

### 4. CEN6-URA3 expression vector

The CEN6-URA GFP dropout backbone vector was constructed by BsaI GG assembly of YTK parts pYTK008 (ConLS’), pYTK047 (GFP dropout), pYTK073 (ConLE’), pYTK074 (URA3), pYTK081 (CEN6/ARS4), pYTK084 (KanR-ColE1 RFP dropout vector).

### 5. An alternate strategy to directly clone annealed sgRNAs in a yeast expression vector

To perform faster direct sgRNA-Cas9 plasmid construction without requiring three independent cloning strategies (as described in ***section 2***), we constructed **pCas9-GFPdo** to directly clone the annealed sgRNA primers into yeast expression vectors. The sgRNA expression unit from the pYTK050 plasmid was amplified via PCR using primers with BsaI sites that generated overhangs similar to ConL1 (forward primer) and ConRE (reverse primer) to clone in CEN6-URA or CEN6-*KanMX* yeast expression vectors. Finally, using the GG reaction protocol, the sgRNA TU PCR was assembled into the CEN6-*URA3* or CEN6-*KanMX* backbone with the Cas9-TU1 vector to create a direct knockout (**pCas9-GFPdo**) vector.

### 6. An alternate strategy to directly clone multiple sgRNAs TUs in a yeast expression vector for multiplexing in MERGE^MX^

The YTK kit allows cloning multiple sgRNA expression cassettes in a yeast expression vector. However, the strategy involves three cloning steps. To build >1 sgRNAs TUs directly into the yeast expression vector without generating intermediate connector vectors, we designed the **pCas9-GFPMX** (CEN6-*URA3*) vector. Primers were designed to PCR amplify the GFP expression cassette (for *E. coli* expression, pYTK047) with GG enzyme sites to clone GFP (BsaI for cloning the GFP insert, the sites are lost after GG cloning). The BsmBI sites, with overhangs similar to ConLS and ConRE, designed in the primers, allow the PCR-based sgRNA TU cloning (obtained by ligating annealed sgRNA primers in pYTK050) in CEN6-*URA3* yeast expression vector (**Table S1**). The primers used to amplify the sgRNA expression units now harbor connector overhangs to clone >1 sgRNA expression units in tandem, eliminating the need to individually make each connector clone.

### 7. Repair design and construction

Repair DNA was designed to be a linear DNA molecule that contained the human coding sequence, from the start codon to stop, with at least 100bp of flanking homology to the yeast genome immediately upstream and downstream of the native yeast start and stop codons (**Figure S3B**). They were constructed by PCR using human ORFeome^37^ or MGC^38^ clones as templates, using primers with long extensions providing the homology sequence. Repair template PCRs were performed with Accuprime Pfx (Thermo) as multiple 100μl reactions according to the manufacturer’s protocol, combined and purified using the Zymo DNA Clean or Qiagen PCR Cleanup and Concentrator −25 kit. Approximately 5μg of repair DNA was included in the transformation mix during sequential replacement transformations.

### 8. Humanization procedure

Yeast transformations were performed using the Frozen-EZ Yeast Transformation II Kit from Zymo Research (Cedarlane). In short, cells were grown to mid-log phase, washed with kit Solution 1, resuspended in kit Solution 2 and either used directly to transform or frozen at −80 C for future use. In general, we transformed approximately 1μg of the KO plasmid along with 5μg of repair DNA, which yielded anywhere from single to dozens of colonies, depending on transformation efficiency and the number of simultaneous molecules necessary. In general, nearly all screened clones were successful. For initial single replacement in the wild-type strains, we tested two sgRNA sequences for each gene. These were tested in parallel, together with a no-repair control transformation to assay for sgRNA effectiveness. Successful sgRNAs show zero or very few colonies in the no-repair control transformation (**CFU_O_/CFU_E_ =∼0, Figure 1A & S4A**).

### 9. Engineering Carotenoid genes at the landing pad loci

Single-sgRNA CRISPR reagents targeting three landing pad loci (*511B*, *USERX*, *FGF20*)^39^ were generated as described in ***section 1***. The repair templates were generated as previously described in ***section 7***. Specifically, carotenoid gene transcription units were amplified from plasmids harboring the transcription units using the primers with 80bp homology to the landing pad loci^40^ (**Table S1**). Each landing pad locus was edited in the SGA strain (Mat A) to harbor a carotenoid gene transcription unit (*511B* for *CrtE*, *USERX* for CrtI, and *FGF20* for CrtYB) driven by strong constitutive yeast promoters. The transformation was carried out using the Zymogen frozen EZ yeast transformation kit (Cedarlane), and colonies were confirmed by PCR using primers specific to the landing pad or the carotenoid gene loci as described below (**Figure S4B** & **Table S1**).

### 10. Clone verification

Clones were initially screened by colony PCR using a rapid DNA isolation method and colony PCR^41^. Forward primers for PCR screening were designed such that the upstream primer would bind in the yeast genome approximately 150bp - 500bp upstream of the yeast ORF or the landing pad loci, to ensure it was outside of the homology region used for repair. Two reverse primers were designed for each locus, one binding into the yeast and the other within the human or carotenoid gene sequence. To facilitate multiplex PCR screening, each pair amplifies different-sized bands for the yeast and human or carotenoid genes. Following plasmid loss, clones were further verified by directly sequencing a PCR product using Sanger sequencing.

### 11. Plasmid loss to perform sequential editing of loci

Successful clones were subjected to a plasmid loss procedure to alleviate any stress incurred by the constitutive expression of the Cas9 protein and allow further knockout plasmids to be transformed. To avoid any potential stress-induced defects, we rarely used 5-FOA counter-selection to force loss of the URA3 plasmid, instead relying on spontaneous plasmid loss. One two methods were used to identify spontaneous plasmid loss. Clones were grown overnight in YPD and then again either spread onto YPD (100μl of 1:1000 dilution) and replica plated on SC-Ura and YPD or patch plated (typically 6-12 colonies) to both minimal media lacking uracil and YPD. This procedure resulted in around 10-60% of colonies losing the plasmid (estimated). For a plasmid harboring URA3 selection, the strains that lost the plasmid were identified using 5-FOA selection.

### 12. Growth curves

Cells were diluted to approximately 0.01-0.02 OD660 (∼2-5 x10^5^ cells) in 150μl YPD across 3-4 replicates. They were grown and read in a BioTek Synergy H1 or Sunrise Tecan microplate reader, with continuous double orbital shaking at 30℃ or 37℃. Reads were taken while stationary every 10-20 minutes, and experiments were run for at least 24-36 hours. For spot assays, strains were grown overnight in YPD at 30℃ and spotted with serial dilutions on YPD agar. The plates were incubated for 2-3 days at 23℃, 30℃ and 37℃.

### 13. CRISPR-based selection (CELECT) to identify and enrich unique genotypes

To test the efficiency of CRISPR selection, a subset of engineered strains (Haploid Mat A) were mixed as a pool. Each strain was cultured overnight in YDP individually and back-diluted into a pool the next day at equal OD and grown to mid-log phase. The strains used in the mix were all of the same mating types to avoid random mating events during the experiment. Competent cells were prepared and transformed with various pCas9-sgRNA^locus^ plasmids. We performed the experiments as biological and technical replicates using different genotypes in the mix. The same competent cell mix was either transformed with pCas9 alone or with various **pCas9-sgRNA*^WT^* ^locus^** vectors, each harboring a different yeast-specific sgRNA expression unit. Competent cells were generated using Zymo Research Frozen EZ kit transformation protocol. Each transformation was plated on SD-URA. pCas9 alone served as a positive control allowing the growth of all genotypes in the mixture. **pCas9-sgRNA*^ADE2^*** was used to visually observe and quantify the efficiency of the selection. The plasmid will eliminate all the other genotypes except the *ade2Δ::kanMX* (red colonies due to the disruption of the *Ade2* gene). pCas9-sgRNA*^RPT5^*was used as a negative control as the yeast *Rpt5* gene is present in all the strains such that yeast cells should survive this transformation. Colonies from each Petri plate were picked randomly and genotyped by locus-specific PCR to confirm the CRISPR reagent exclusively selected a unique genotype as described in section ***10***. (**Figure 1C**).

### 14. Mating and MERGE^0^ assays

The mating assays were carried out in the following ways:

***14.1.*** The engineered strains with single gene modification in BY4741 (Haploid Mat A) backgrounds were mated with SGA-haploid strains harboring haploid specific markers (SGA Mat A; *can1Δ::STE2pr_Sp_his5his3Δ1leu2Δ0ura3Δ0;* select on SD-His+Canavanine & SGA Mat α; *lyp1Δ::STE3pr_LEU2 his3Δ1leu2Δ0ura3Δ0;* select on SD-Leu+Thialysine). Each parental strain was transformed with two different empty vector plasmids with distinct selectable markers *URA3* and *KanMX*, and mated diploids were selected on SD-Ura+G418. The haploid strains were grown overnight in 5ml selection media (SC-Ura or YPD +G418). The strains were mixed in a rich medium and incubated at 30°C for 3-4 hrs. with shaking. 500µl of the mated mixture was washed with distilled water, and 10µl of the mix was plated at different dilutions on solid agar media using SD-URA+G418 selection and incubated for 2-3 days at 30°C. Plasmids were cured using a previously described strategy in section ***11***. The heterozygous colonies were confirmed by PCR genotyping using primers specific to the wild-type and engineered loci as described in section ***10***. The confirmed heterozygous strains for all proteasome and carotenoid strains were transformed with CRISPR plasmids targeting wild-type loci and converting the parent heterozygous strain into homozygous diploid strain. Colonies were picked randomly from the plate and genotyped for homozygosity using the same set of primers, confirming the loss of yeast and the presence of humanized or carotenoid loci.

***14.2.*** Alternatively, instead of an empty vector, the engineered humanized or carotenoid strains were transformed with a CRISPR reagent for which the corresponding strain harbors a CRISPR-resistant locus. The subsequent mating with an opposite mating-type strain with an empty vector (pCas9 alone or a backbone plasmid) results in the loss of heterozygosity and conversion of single loci (**MERGE^0^**). In each scenario, the number of colonies obtained on plates transformed with CRISPR reagent (**CFU_O_**) were compared with empty vector transformations (**CFU_E_**) to calculate the efficiency of **MERGE^0^**. Each experiment was performed at least 3 times.

### 15. Sporulation of diploid and selection of haploid yeast strains

Diploid strains were sporulated in a media containing 0.1% potassium acetate and 0.005% zinc acetate for 4-7 days. For Tetrad dissection, spores were spun down at 5000rpm for 5 minutes, resuspended in 200μl of 20mg/ml Zymolyase and incubated at 37°C for 25-30 minutes. The mix was then incubated at −20°C to stop the reaction. Cells were thawed on ice, and 20ul of the mix was plated into YDP for Tetrad dissection. Tetrad Dissection was performed using Spore play (Singer Instruments for every engineered proteasome spore on YPD and then replicated on SGA selection. For carotenoid and humanized loci in SGA background strain, the sporulation mix was directly plated on SGA selection. Locus-specific PCR verified the haploid strains for the presence or absence of engineered loci.

### 16. Estimating the efficiency of MERGE^0^ using yeast ADE2 locus as a readout

The yeast *ADE2* locus was used to estimate **MERGE^0^** using a CRISPR-resistant allele *ade2Δ::kanMX* on the homologous chromosome as a repair template. The sgRNAs were designed using Geneious version 11 as described in ***section 1*** (**Table S1**). The **pCas9-sgRNA*^ADE2^*** vector showed ON-target activity causing lethality in wild-type haploid or diploid strains. Occasional colonies show a red colony phenotype suggesting mutations via NHEJ at the *ADE2* locus. To estimate **MERGE^0^** efficiency, the **pCas9-sgRNA*^ADE2^*** was transformed in a diploid hetKO *ADE2*/*ade2Δ::kanMX* obtained from the yeast “Magic Marker’’ hetKO collection^26^. To further verify that the red colonies observed after **MERGE^0^** are not due to the mutations in the *ADE2* locus (NHEJ) and instead due to conversion to *ade2Δ::kanMX* locus (HDR), the heterozygous diploid (ADE2/*ade2Δ::kanMX*) strain transformed with a control (**pCas9 alone**) and **pCas9-sgRNA*^ADE2^*** plasmid were sporulated followed by tetrad dissection (**Figure S5A**). Haploid spores were selected on YPD or YPD + G418 (200µg/ml). Spores harboring a control plasmid showed the expected 2 : 2 (red : white phenotype) and 2 : 0 (G418 resistant, G418 sensitive) phenotype compared to the **pCas9-sgRNA*^ADE2^*** transformed cells that show 4 : 0 (red, white) colonies and 4 : 0 (G418 resistant, G418 sensitive) colonies phenotype, respectively, suggesting the conversion to *ade2Δ::kanMX* rather than the mutation of *Ade2* locus. In contrast, the co-transformation of **pCas9-sgRNA*^ADE2^*** and oligo as a repair template harboring (100bp homology, 5X molar excess than the plasmid) to the 5’ and 3’ UTRs of *ADE2* locus in haploid wild-type yeast cells, shows resulted in significantly fewer survivors (**Figure 2C**; % **CFU_O_**/**CFU_E_**= 21.6+/-SD; N=4). However, this method is still far less-efficient than **MERGE^0^**(**Figure 2C**; ∼100% vs 21.6%).

### 17. Using MERGE^0^ to perform one-step gene essentiality assays in yeast

HetKO diploid strains for 7 *α* proteasome core genes were obtained from the yeast “Magic Marker’’ hetKO collection^26^. The strains were transformed with either a single-sgRNA CRISPR reagent targeting the corresponding yeast *α* proteasome genes or the empty vector control and selected on SD-URA+G418. CRISPR reagent transformed plates caused lethality in 6 of 7 *α* proteasome hetKO strains, except in the case of non-essential *α3*. A similar assay in the hetKO *ADE2*/*ade2Δ::kanMX* strain showed viable homozygous null cells for a non-essential *ADE2* locus.

### 18. Testing MERGE^0^ efficiency for all yeast loci on Chromosome I

All the strains harboring heterozygous knockout diploid loci from the yeast “Magic Marker” collection on chromosome I were arrayed in a 96-well format^26^ (**Figure S6A**). A2 is the locus close to the left telomere, F1 near the centromere, and G10 close to the right telomere (**Figures 2D & S6A**). The sgRNA targeting the *KanMX* cassette was designed using the strategy mentioned earlier (**Table S1**). Since the bacterial selection for the previously described (***section 4***) yeast shuttle vector harbors a *Kan^R^* cassette identical to the *KanMX* cassette in the case of hetKO strains, we designed a new expression vector with *AmpR* selection. The plasmid showed a lethal phenotype in the strains harboring the *KanMX* allele (haploid or homozygous diploid) in the genome (**Figure 2D**). Approximately 1µg of **pCas-sgRNA*^KanMX^*** was transformed for each transformation.

First, we performed a pilot assay using 5 hetKO strains with genes located at various regions on chromosome I (**Figure S6B**). These strains showed no lethality (**CFU_o_/CFU_E_ of ∼1**). Each strain was inoculated in 800µl of YDP + His(50 mg/L) + G418 (200 µg/ml) overnight in 96-well format. The following day, cultures were back-diluted and grown to the mid-log phase. Competent cells were generated in a 96-well format using the Gietz yeast transformation protocol (Gietz, 2007). Each transformation mix was transformed with either 1µg of pCas9 alone or 1µg of the **pCas9-sgRNA*^KanMx^*** plasmid. An equal amount of the transformed cells were plated onto SD-URA and SD-URA + G418. All of the yeast loci (except for A10- *CNE1*) on chromosome I show ***MERGE^0^*** mediated conversion of *KanMX* to wild-type yeast locus irrespective of the position of the gene along the chromosome (**CFU_O_/CFU_E_ =∼1**) and simultaneously lost the KanMX cassette (**Figure S6A**).

### 19. Performing MERGE^1^ and MERGE^MX^ at 2 or more loci using humanized proteasome and carotenoid strains

CRISPR plasmids expressing 2 or more sgRNA cassettes in tandem were designed and constructed as described in sections ***1 - 6***. Each single-humanized α proteasome (BY4741, Mat A) was mated with SGA strain (Mat α) as described in section ***14***. Heterozygous strains from each mating mix were confirmed by locus-specific PCR genotyping as in section ***10***. ***MERGE^0^*** enabled the conversion to humanized loci. The next round of mating in SGA strain (Mat A) followed by **MERGE^0^** generated humanized strains in both mating-type SGA backgrounds. Carotenoid strains were directly generated using SGA compatible strains described in **section *9,*** followed by **MERGE^0^** to obtain strains in both mating-types.

Each double-sgRNA CRISPR reagent was lethal in singly engineered strains (**Figure S7B**). Mating between each distinct single-engineered genotype combines the loci as heterozygotes. Transformation of 0.5-1µg of double-sgRNA CRISPR reagent in heterozygous diploid strains either showed no lethality (viable genotype, **CFU_O_/CFU_E_ =∼1**) or a lethal phenotype (inviable genotype, **CFU_O_/CFU_E_ =∼0**) (**Figures 3B, 3C &S7C**). Homozygosity was confirmed by randomly picking colonies after **MERGE^1^** and performing a locus-specific PCR confirming the presence or absence of yeast or engineered genotype.

For ***MERGE^MX^*** to simultaneously target >2 loci, the CRISPR reagents were generated as described in sections ***2-4*** and ***6***. The haploid strains harboring combined engineering loci were mated with either a wild-type or strains carrying a different set of engineered loci. The diploids were transformed with either an empty vector or a multiplex-sgRNA CRISPR reagent. The genotypes were confirmed by locus-specific PCR as described in section ***10***.

### 20. Performing MERGE^M&M^ to combine 2 loci from the mated mix of single-humanized proteasome genes

Single-humanized α proteasome strains *Hsα1*, *Hsα3*, and *Hsα7* of both mating-types were allowed to mate randomly in a mix. Each strain was inoculated at 0.3 OD in 2 ml YPD overnight, then added to a mix at equal OD and incubated on a shaking incubator for 4-6 hrs 30°C. 500µl of the mix was further cultured overnight at starting 0.3 OD, followed by competent cell preparation. The mixture was transformed with either an empty vector or double-sgRNA CRISPR reagent. Several colonies were observed post ***MERGE^M&M^*** for *Hsα1α7,* and Hs*α3α7* paired genotypes. PCR-based genotyping confirmed the combination as a homozygous double-humanized diploid.

### 21. Testing the potential of CRISPR reagent to target multiple loci using the Green Monster strain

To address the scalability of targeting multiple yeast loci, we used the Green Monster (GM) strain^17^. The sgRNA target sequence was designed in GFP (S65T) variant using Geneious version 11 as described in section ***1***. **pCas9-sgRNA*^GFP^*** was constructed using the method of direct cloning strategy as described in section ***5*** using a backbone with *Amp^R^* (*E. coli*) and *KanMX* (yeast) selection. The Mat α mating-type Green Monster was used *MATα lyp1Δ his3Δ1 leu2Δ0 ura3Δ0 met15Δ0 can1Δ::GMToolkit-α [CMVpr-rtTA NatMX4 STE3pr-LEU2]*. The OFF-target activity of **pCas9-sgRNA*^GFP^*** was tested in a haploid WT strain with no *GFP* gene showing a **CFUo/CFU_E_ =∼ 1**. The ON-target activity was measured transformation in natively tagged C-terminal fusion of GFP to *BRO1* gene *(Bro1-GFP)* obtained from the yeast GFP collection^42^.

To estimate the loss-of-function mutations at the GFP locus, cells from an empty vector and **pCas9-sgRNA*^GFP^*** transformed GM yeast cells were subjected to flow cytometry (BD Accuri™ C6 Plus Flow Cytometer). Cells were grown in 96-well format in YDP overnight and back diluted in SC media with 10µg/ml of doxycycline for ∼48 hrs. The culture was diluted 1 in 10 in water and passed through the flow cytometer. Using BY4741 yeast with no GFP as a control, the cells showed mostly background fluorescence as expected. GM cells transformed with an empty vector show high fluorescence as this strain harbors 16 GFP cassettes serving as a positive control. Finally, random colonies were picked and pooled from haploid GM cells transformed with **pCas-sgRNA^GFP,^** and the mixture passed through the flow cytometer. The cells show little GFP expression similar to the wild type, suggesting ON-target NHEJ-mediated mutations at nearly all GFP loci (**Figures S8**).

### 22. *Performing MERGE^MX^* to assemble an entire Carotenoid pathway in yeast using mating and sporulation cycles

SGA strains of both mating-types harboring single-carotenoid transcription units at the landing pad loci were generated in sections ***9 and 14***. The strains were cultured independently overnight, followed by mixing at a similar 0.3 OD and cultured on a shaking incubator for 5-6 hours. A hole was made on the lid of the Eppendorf tube to allow aeration, and the mixture was incubated at 30℃ for 12-16 hrs. The formation of diploids was confirmed by light microscopy. Next, 500µl of culture was centrifuged at 3500 rpm for 5 minutes. The supernatant was removed and resuspended in 2 - 5ml of sporulation media, followed by incubation on a rotating shaker for 5-8 days at room temperature. Next, 500 ul of sporulation mix was centrifuged at 3500 rpm for 5 min and treated with Zymolyase. After the first sexual cycle, the cells were centrifuged, washed with water, and resuspended in rich media YPD. The cycle was repeated to generate a second set of diploids. The mixture was incubated for 4-6 hours in a shaking incubator and incubated at 30℃ for 2 days. The confirmation and appearance of most diploids were again confirmed with a light microscope, and the 100µl of culture was inoculated overnight to make competent cells. The competent cells were transformed with either an empty vector or a triple-sgRNA CRISPR reagent **pCas9-sgRNA**^511^***^B,USERX,FGF20^***. For the most part, we followed a GFP monster protocol. However, we did not use haploid or diploid-specific selection^43^. Instead, CRISPR selection was sufficient to enrich unique combined genotypes.

### 23. Genomic DNA isolation & Whole Genome sequencing, and SNP analysis

Genomic DNA was purified from single*-*humanized *Hsα1* and quintuple-humanized *Hsα1,α2,α3,α4,α7* strains using the Monarch Nucleic acid purification kit according to the manufacturer’s protocol (NEB). Spheroplasts were obtained before the genomic DNA extraction for a high-quality DNA prep. Wild type and engineered strains were sequenced using Illumina MiSeq 2×150 at 30x coverage using 150-bp paired-end reads. The Geneious Pro Software [PMID: 22543367] and its included tools were utilized for pairing paired-end sequences, trimming ends and adapters based on quality using the BBDuk tool, and reference mapping using the Geneious Read Mapper algorithm at medium-low sensitivity, iterated 5 times. Reads were mapped to a reference BY4741 strain (for the engineered strain, the same reference sequence was used, but replacing the sequences of the 5 engineered genes with their human ortholog to ensure alignment with the humanized loci reads) using the Geneious Prime Software 2020.2.4 algorithm. Mapping was run at medium sensitivity: word length to allow for matching was set to 18 with a maximum permitted mismatch of 20% of the read length, maximum mismatches per read were set to 20%, a minimum 80% overlap was required, and reads with errors were set to accurately be mapped to repeat regions - this was iterated 5 times to give the final mapping. For analysis, low coverage regions of below 2 standard deviations from the mean were excluded from SNP-calling, and only SNPs with a variant frequency of 0.90 or higher were considered. SNPs were called with a minimum variant frequency of 0.25, and low coverage regions below 2 standard deviations from the mean were excluded. SNPs that were unique to the engineered strain (i.e. not in the mappings of the wild-type strain) were marked. To ensure SNPs were not introduced by off-target CRISPR effects, 300bp up- and down-stream of each SNP was searched for CRISPR-gRNA alignment at 75% sequence similarity.

